# STRIPAK directs PP2A activity toward MAP4K4 to promote oncogenic transformation

**DOI:** 10.1101/823096

**Authors:** Jong Wook Kim, Christian Berrios, Miju Kim, Amy E. Schade, Guillaume Adelmant, Huwate Yeerna, Emily Damato, Amanda Balboni Iniguez, Selene K. Swanson, Laurence Florens, Michael P. Washburn, Kim Stegmaier, Nathaniel S. Gray, Pablo Tamayo, Ole Gjoerup, Jarrod A. Marto, James A. DeCaprio, William C. Hahn

## Abstract

Alterations involving serine-threonine phosphatase PP2A subunits occur in a range of human cancers and partial loss of PP2A function contributes to cell transformation. Displacement of regulatory B subunits by the SV40 Small T antigen (ST) or mutation/deletion of PP2A subunits alters the abundance and types of PP2A complexes in cells, leading to transformation. Here we show that ST not only displaces common PP2A B subunits but also promotes A-C subunit interactions with alternative B subunits (B’’’, striatins) that are components of the Striatin-interacting phosphatase and kinase (STRIPAK) complex. We found that STRN4, a member of STRIPAK, is associated with ST and is required for ST-PP2A-induced cell transformation. ST recruitment of STRIPAK facilitates PP2A-mediated dephosphorylation of MAP4K4 and induces cell transformation through the activation of the Hippo pathway effector YAP1. These observations identify an unanticipated role of MAP4K4 in transformation and show that the STRIPAK complex regulates PP2A specificity and activity.

## Introduction

Protein phosphorylation plays a regulatory role in nearly all biological processes and dysregulation of protein phosphorylation contributes to many diseases. In cancer, both kinases and phosphatases have been implicated in the pathogenesis of specific cancers and several small molecule kinase inhibitors are standard treatments in such cancers. In addition, several phosphatases have been identified as tumor suppressors (Sablina et al. 2007, Lawrence et al. 2014).

PP2A, an abundant serine/threonine phosphatase in mammalian cells, is comprised of three subunits: A (structural), B (regulatory), and C (catalytic). The A and C subunits form the core enzyme and interact with different B regulatory subunits to create many distinct PP2A enzymes (Pallas et al. 1990, Chen et al. 2007, Cho et al. 2007, Shi 2009, Sents et al. 2013). Moreover, there are two A, two C isoforms, and at least 4 classes of B subunits B, B’, B’’, and B’’’ (striatins), each of which exist as several different isoforms. Although the prevailing view is that the B subunits provide substrate specificity, how B subunits accomplish this regulation remains unclear (Shi 2009, Hertz et al. 2016).

Genome characterization studies of human cancers have identified recurrent mutations and deletions involving PP2A subunits. Indeed, the PP2A Aα (PPP2R1A) subunit ranks among the most recurrently mutated gene across many cancer types (Lawrence et al. 2014). Notably, mutations in Aα occur at high frequency in premalignant endometrial lesions (Anglesio et al. 2017). PP2A is also a target of the Small T antigens (ST) of SV40 and other polyomaviruses including the human oncogenic Merkel cell polyomavirus (Pallas et al. 1990, Chen et al. 2004, Cheng et al. 2017) and this interaction contributes to cell transformation (Hahn et al. 2002). Structural studies have shown that ST disrupts formation of a functional PP2A holoenzyme by displacing or hindering B subunit access to the PP2A core-enzyme (Chen et al. 2007, Cho et al. 2007). However, it was also shown that ST has a lower binding affinity in vitro for the PP2A core enzyme than B’ subunits, which suggests that ST interaction with the core enzyme may either occur prior to the B subunit binding, or ST directly inhibits PP2A activity independently of subunit assembly (Chen et al. 2007).

Several investigators have used mass spectrometry to identify proteins that interact with PP2A (Goudreault et al. 2009, Herzog et al. 2012). These studies identified a large complex called the Striatin-interacting phosphatase and kinase (STRIPAK) complex (Goudreault et al. 2009). The STRIPAK complex contains Striatin family (STRN) proteins, several kinases, scaffolding proteins, and PP2A subunits. Indeed, striatins were initially described as non-canonical PP2A regulatory subunits (B’’’ subunits) (Moreno et al. 2000). STRIPAK complexes have also been shown to associate with members of the GCKIII kinase subfamily (MST3, STK24, and STK25) (Kean et al. 2011). In addition, mitogen-activated protein kinase kinase kinase kinase 4 (MAP4K4), a Ste20-like kinase, although not an obligate member of the STRIPAK complex, associates with STRIPAK (Frost et al. 2012, Herzog et al. 2012, Hyodo et al. 2012). We also identified members of the STRIPAK complex, including STRN3, STRN4, STRIP1, and MAP4K4 in complex with SV40 ST (Rozenblatt-Rosen et al. 2012). Although STRIPAK comprises multiple signaling enzymes, it is unclear how disruptions to the biochemical complex integrate with or disrupt phosphorylation cascades; or whether these signaling alterations synergize with ST to mediate cellular transformation.

MAP4K4 is a serine/threonine kinase that was initially found to activate the c-Jun N-terminal kinase (JNK) signaling pathway (Yao et al. 1999), downstream of TNF-α. MAP4K4 has also been implicated in a large number of biological processes including insulin resistance, focal adhesion disassembly, as well as cellular invasion and migration (Collins et al. 2006, Tang et al. 2006, Yue et al. 2014, Danai et al. 2015, Vitorino et al. 2015). Recent studies have shown that MAP4K4 phosphorylates LATS1/2, activating the Hippo tumor suppressor pathway, leading to YAP1 inactivation (Mohseni et al. 2014, Meng et al. 2015, Zheng et al. 2015). Here, we investigated the role of the STRIPAK complex and MAP4K4 in human cell transformation driven by SV40 ST and found that kinase inactivation or partial suppression of MAP4K4 replaces expression of ST in the transformation of human cells.

## Results

### Identification of MAP4K4 as a candidate phosphoprotein targeted in cells transformed by PP2A perturbation

Human embryonic kidney (HEK) epithelial cells expressing SV40 Large T antigen (LT), the telomerase catalytic subunit (hTERT), and oncogenic HRAS (referred to as HEK TER hereafter) have served as a useful model system to identify pathways and protein complexes that can functionally substitute for SV40 ST in promoting transformation, including partial depletion of PP2A (Chen et al. 2004, Sablina et al. 2010). These cells, upon expression of SV40 ST or partial knockdown of PP2A Aα or Cα subunits, become tumorigenic (Hahn et al. 2002, Chen et al. 2004). Prior studies have shown that expression of ST, or partial inhibition of certain PP2A subunits, causes increased phosphorylation of PP2A substrates (Sablina et al. 2008, Sablina et al. 2010).

To assess the serine/threonine phosphorylation events that are associated with transformation induced by ST or by partial knockdown of PP2A, we performed global Isobaric Tags for Relative and Absolute Quantitation (iTRAQ) phosphoproteomic profiling of HEK TER cells expressing ST (HEK TER ST) or in which expression of the PP2A Aα, Cα, or B56γ subunits was suppressed using previously characterized shRNAs (Fig. 1A) (Sablina et al. 2010). We also confirmed by anchorage-independent (AI) growth assays that these genetic perturbations promoted the transformation phenotype as previously described (Fig. S1A) (Sablina et al. 2010). After mass spectrometry analysis of the phosphopeptides altered across these conditions, we identified 6025 phosphopeptides corresponding to 2428 individual proteins reproducibly detected in two replicate experiments. After processing and normalization of the raw data (see methods for details), we performed comparative marker selection analysis (Gould et al. 2006) to identify candidate phosphoproteins that were most significantly correlated with the transformation phenotype (Fig. 1A). In consonance with previous studies (Ratcliffe et al. 2000, Kuo et al. 2008), we observed an increase in phosphorylation of direct or indirect targets of PP2A, including AKT1S and β-catenin (CTNNB1) in cells which were transformed by either expressing ST or partial knockdown of PP2A Cα subunit in HEK TER cells (Fig. 1B) (Sablina et al. 2010). Conversely, we also observed decreased phosphorylation on multiple proteins in cells transformed by ST or by PP2A perturbation (B56γ1, Cα2). Notably, the phosphorylation signature for transformation included four distinct sites on MAP4K4 (T804, S888, S889, S1272, p<0.05, Fig. 1B).

**Figure 1.**
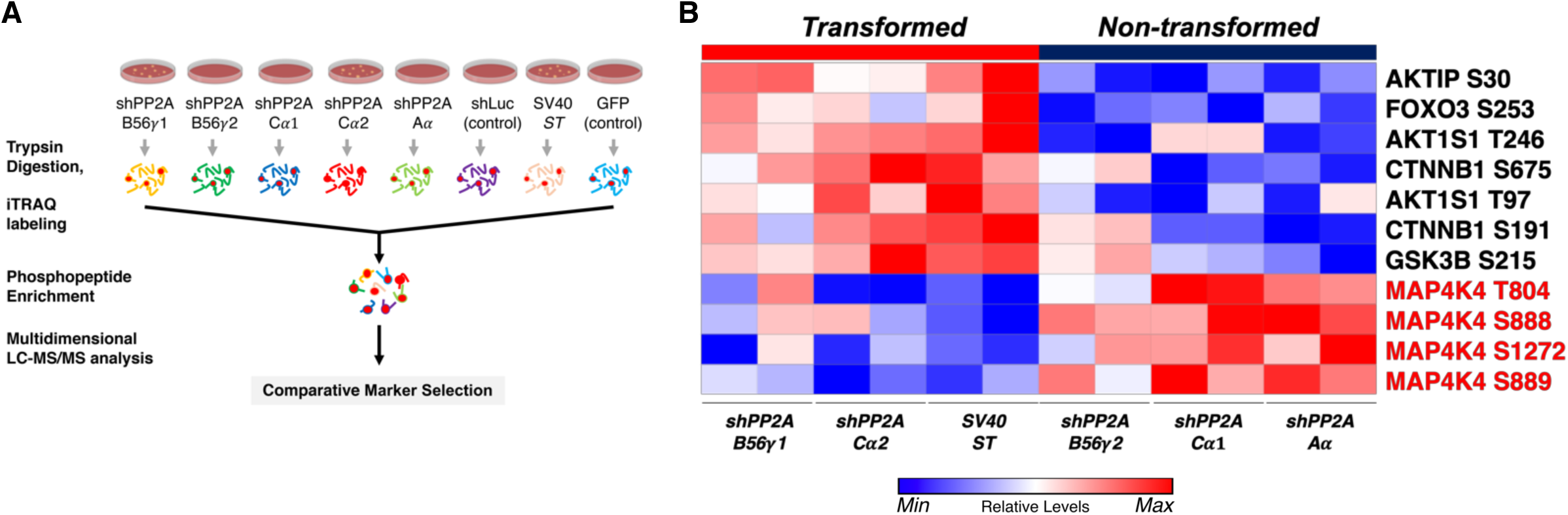
Global phosphoproteomic analysis identifies MAP4K4 dephosphorylation in cells transformed by PP2A perturbation. (A) Schematic illustrating the global phosphoproteomics experiment. (B) The heatmap depicts phosphopeptides that are either positively or negatively correlated with the transformation phenotype (p<0.05, FDR<1). Each column represents individual samples (normalized to shLuc for shPP2A or in the case of ST to GFP control). The sample designations after the normalization and comparative marker selection analysis are shown below the heatmap, with each sample shown in replicates. A selected subset of phosphorylated sites which distinguishes transforming and non-transforming perturbations are shown.

Our previous systematic analysis of SV40 ST identified MAP4K4, in addition to PP2A and other STRIPAK components in the same complex (STRN4, STRN3, CTTNBP2NL, FAM40A, MAP2K3, STK24, PPP2R1A) (Rozenblatt-Rosen et al. 2012). To confirm these interactions, we generated lentiviral C-terminal Flag-HA Tandem Affinity Purification (CTAP) constructs for SV40 ST, as well as ST from 3 closely related Human Polyoma Viruses (HPyV) including JCPyV-CY, JCPyV-Mad1, and BKPyV, along with GFP as a negative control. We introduced these viral proteins into HCT116 cells and performed HA-tag immunoprecipitations (IP) from lysates of cells expressing ST from the respective viruses. We confirmed co-complex formation between SV40 ST and MAP4K4, as well as with STRIPAK components PP2A Cα, STRN3 and STRIP1 (Fig. S1B). We observed that ST of JCPyV and BKPyV, the two most closely related HPyVs to SV40, also interacted with STRN3, STRIP1 and PP2A Cα but not MAP4K4, indicating that the interaction of SV40 ST and MAP4K4 was unique to SV40 ST. The association of ST with B’’’ subunits (striatins) was unexpected, because ST was previously reported to primarily bind PP2A Aα and displace most B subunits (Pallas et al. 1990, Chen et al. 2007, Cho et al. 2007, Sablina et al. 2010). These results raised the possibility that ST modulates MAP4K4 phosphorylation via PP2A activity associated with the STRIPAK complex.

### Partial knockdown of MAP4K4 promotes cell transformation

To determine if MAP4K4 and other SV40 ST interacting proteins participated in cell transformation, we created and stably expressed two distinct shRNAs targeting each of several SV40 ST interacting proteins including STRN3, STRN4, STRIP1, MARCKS, MAP4K4, and STK24 in HEK TER cells. We then assessed the ability of each of these shRNAs to promote AI growth, a readout for the transformed phenotype (Fig. 2A, S2A). As expected, expression of SV40 ST or partial knockdown of PP2A Cα subunit in HEK TER cells induced robust AI growth (Fig. S2A). Among the STRIPAK components, we found that one of the two shRNAs targeting MAP4K4 (shMAP4K4-82) elicited a potent transformation phenotype (Fig. 2B-C, S2A). To ensure that the observed phenotype was specific to targeting MAP4K4 and not due to an off-target effect of this shRNA, we repeated the AI growth assay using 8 different MAP4K4-targeting shRNAs including the 2 shRNAs used in the initial experiment (Fig. S2B). In addition, we found that the three shRNAs which promoted HEK TER cells to grow in an AI manner (shRNA-82, 92, 93) only partially suppressed MAP4K4 levels (Fig. S2B). Specifically, we focused on shMAP4K4-82, which promoted the most robust AI growth and knocked down MAP4K4 mRNA levels by 50% (Fig. S2B-C). In contrast, none of the shRNAs that induced more than 50% knockdown of MAP4K4 expression resulted in AI growth (Fig. S2B). This relationship between partial knockdown and cell transformation is similar to what has been reported for the knockdown of PP2A Aα and Cα subunits (Chen et al. 2005, Sablina et al. 2010). To further confirm these data *in vivo*, we performed xenograft experiments to assess tumor formation by subcutaneous injection of immunodeficient mice. Consistent with the *in vitro* studies, shMAP4K4-82 induced potent tumor formation when compared to the shLuciferase (shLuc) control (Fig. 2D-E). These observations suggest that partial knockdown, but not full depletion, of MAP4K4 promotes both transformation and tumor formation.

**Figure 2.**
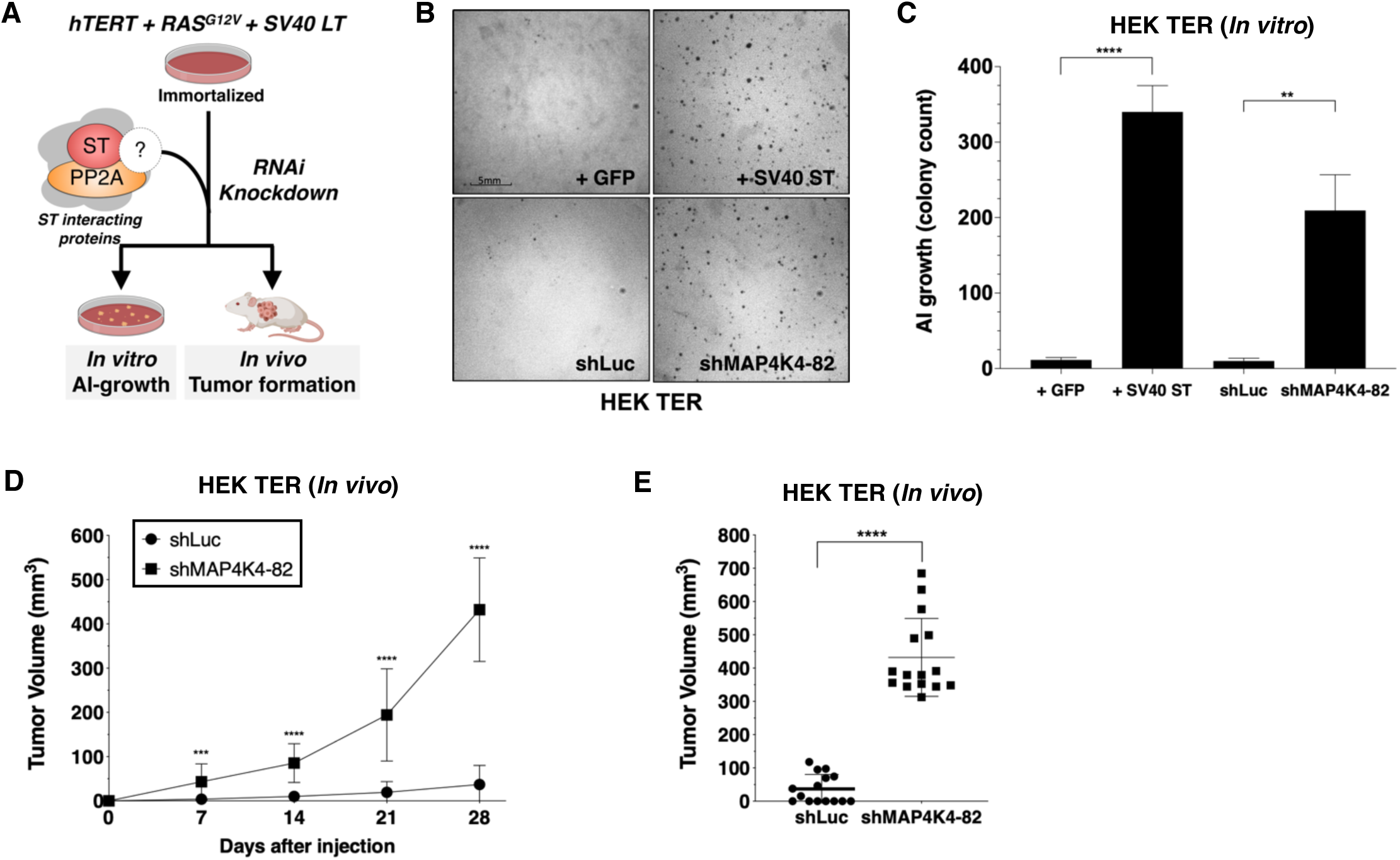
Partial knockdown of MAP4K4 expression promotes oncogenic transformation and tumor formation. (A) Schematic of experimental design to reveal binding proteins that when depleted can substitute for ST in transformation (B) Representative image of AI growth induced by ST or MAP4K4 partial knockdown (Grid shows 5mm). (C) Quantification of AI growth following expression of MAP4K4 shRNA-82 (shMAP4K4), SV40 ST or corresponding controls (GFP or shLuc). Graph depicting tumor volume as a function of time (D) or at endpoint at day 28 (E) for subcutaneous xenografts expressing shLuc control or shMAP4K4-82 in HEK TER cells (Student’s t-test, **p<0.001,***p<0.0001, ****p<0.00001).

### SV40 ST promotes the interaction of MAP4K4 with STRIPAK

To understand the mechanism by which the ST/MAP4K4 axis contributes to cell transformation, we first assessed changes in interactions between MAP4K4 and its binding partners upon ST expression. Specifically, we stably expressed NTAP-MAP4K4 in HEK TER cells expressing either ST or GFP as a negative control. We used Stable Isotope Labeling with Amino Acids (SILAC) to encode proteins in each condition (Fig. 3A). We found that MAP4K4 interacted with STRIPAK components, including STRIP1, STRN3, STRN4 and the PP2A Aα subunit. The interactions between MAP4K4 and the STRIPAK components were increased by 3-4-fold in cells expressing ST relative to the GFP control (Fig. 3B). We tested a series of ST mutants (R21A, W147A, F148A, P132A) that are unable to bind to PP2A Aα (Cho et al. 2007) in 293T cells, and found that these mutant ST proteins were unable to interact with STRN3, a core component of the STRIPAK complex (Fig. S3A), demonstrating that this interaction is dependent on ST binding to PP2A Aα subunit. These observations indicated that MAP4K4 interaction with STRIPAK is enhanced in cells expressing SV40 ST.

**Figure 3.**
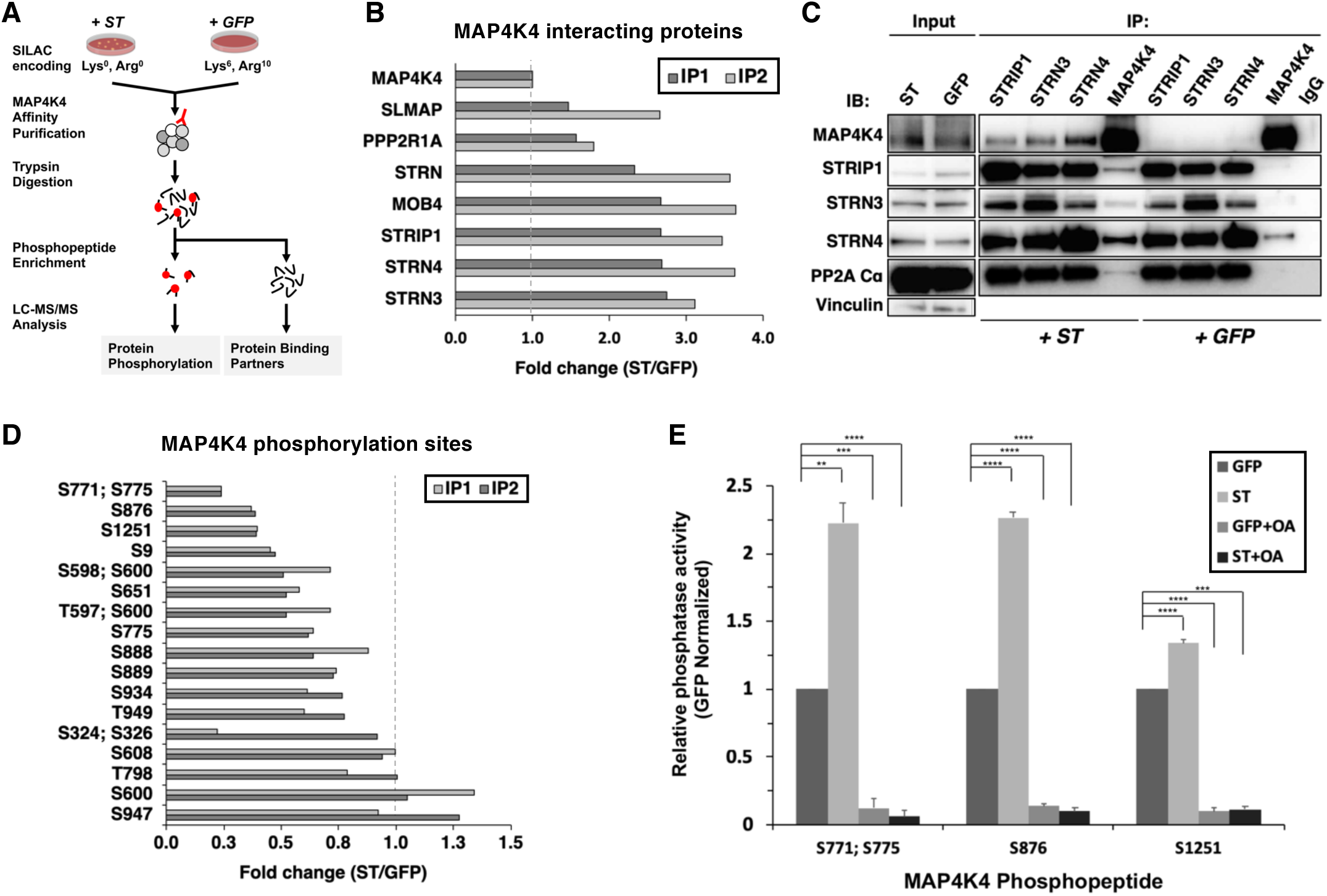
ST promotes MAP4K4 interactions with STRIPAK and MAP4K4 dephosphorylation. (A) Schematic of targeted proteomic analysis of MAP4K4 phosphorylation and interacting proteins in the presence of ST or GFP control. (B) SILAC experiment in which MAP4K4-associated proteins were assessed in cells expressing ST or a GFP control in two biological replicate SILAC experiments (IP1, IP2). The proteins that showed a fold change above 1 have increased interactions with MAP4K4 in ST expressing cells relative to GFP. All values of the retrieved peptides were normalized to the total number of MAP4K4 peptides, prior to calculating the ratios between ST-versus GFP-expressing cells to account for variations in the amount of MAP4K4 after affinity purification. (C) Immunoblot showing results of a Co-Immunoprecipitation (Co-IP) analysis of components of STRIPAK with MAP4K4 in ST- or GFP-expressing cells. ST induced the association of MAP4K4 with STRIPAK components. (D) Quantification of fold changes in the abundance of MAP4K4 phosphorylation across indicated sites (y-axis) in cells expressing ST relative to the GFP control in two independent experiments (IP1, IP2). The phosphosites with fold changes below 1 show a decrease of phosphorylation of MAP4K4 in ST expressing cells relative to GFP. All values of the retrieved peptides were normalized to the total number of MAP4K4 peptides, prior to calculating the ratios between ST-versus GFP-expressing cells to account for variations in the amount of MAP4K4 after affinity purification. (E) After immunoprecipitation of STRN4 from ST-versus GFP-expressing cells, *in vitro* PP2A activity was measured with synthetic MAP4K4 peptides (S771;S775, S876, or S1251) identified in the targeted phosphoproteomic experiments (x-axis). Relative phosphatase activity in ST-relative to GFP-expressing HEK TER cells is shown for each phosphopeptide (y-axis). Okadaic acid (OA) was used to inhibit PP2A activity in parallel conditions (Student’s t-test, **p<0.001, ***p<0.0001, ****p<0.00001).

To corroborate these observations, we performed IP of endogenous STRN3, STRN4, STRIP1, and MAP4K4 and compared the interactions of components of the STRIPAK complex with MAP4K4 in HEK TER cells expressing either ST or GFP. Consistent with the proteomic results, we observed that the interaction of MAP4K4 with the STRIPAK complex was significantly enhanced in the presence of ST (Fig. 3C). We also performed these experiments in normal human fibroblasts (IMR90) expressing ST or GFP (negative control) and confirmed the enhanced binding of MAP4K4 to STRN4 and STRIP1 (Fig. S3B). These observations indicate that interactions between MAP4K4 and STRIPAK components, including STRIP1, STRN3 and STRN4 are enhanced in the presence of SV40 ST.

We next analyzed the enriched phosphopeptides from affinity purified MAP4K4 (Fig. 3A) to better interrogate the full phosphorylation landscape on the kinase. In two independent experiments, we quantified 17 MAP4K4 phosphorylation sites (Fig. 3D). The majority of these sites exhibited reduced phosphorylation in cells expressing ST. These findings further demonstrate that ST mediates dephosphorylation of several distinct MAP4K4 sites.

To evaluate if MAP4K4 dephosphorylation is mediated by the STRIPAK complex, we isolated STRN4 from cells expressing ST or GFP and measured PP2A-specific dephosphorylation activity using synthetic phosphopeptides encompassing MAP4K4 sites S771;S775, S876, or S1251. These sites were selected, because they exhibited the largest change in phosphorylation upon ST expression (Fig. 3D). As a control, we treated parallel samples with okadaic acid (OA), a potent and specific PP2A inhibitor. As expected, we observed that OA treatment eliminated phosphatase activity under all conditions (Fig. 3E). In contrast, co-incubation of MAP4K4 phosphopeptides with STRN4 immune complexes from ST-expressing cells led to dephosphorylation of the S771/S775 and S876 phosphopeptides by greater than 2-fold compared to GFP control, while we found a modest but reproducible increase of dephosphorylation of the S1251 site (Fig. 3E). These observations suggest that ST promotes PP2A-mediated dephosphorylation of MAP4K4 in the STRN4 complex.

### Attenuation of MAP4K4 kinase activity is associated with transformation

Since MAP4K4 phosphorylation at several sites was substantially attenuated in the presence of ST, we assessed whether this decrease in MAP4K4 phosphorylation affected MAP4K4 activity by performing an *in vitro* kinase assay using tandem-affinity purified MAP4K4 from cells that expressed ST or GFP control. We found that the activity of MAP4K4 was reduced in ST-expressing cells compared to cells that expressed GFP control (Fig. 4A, Fig. S3C).

**Figure 4.**
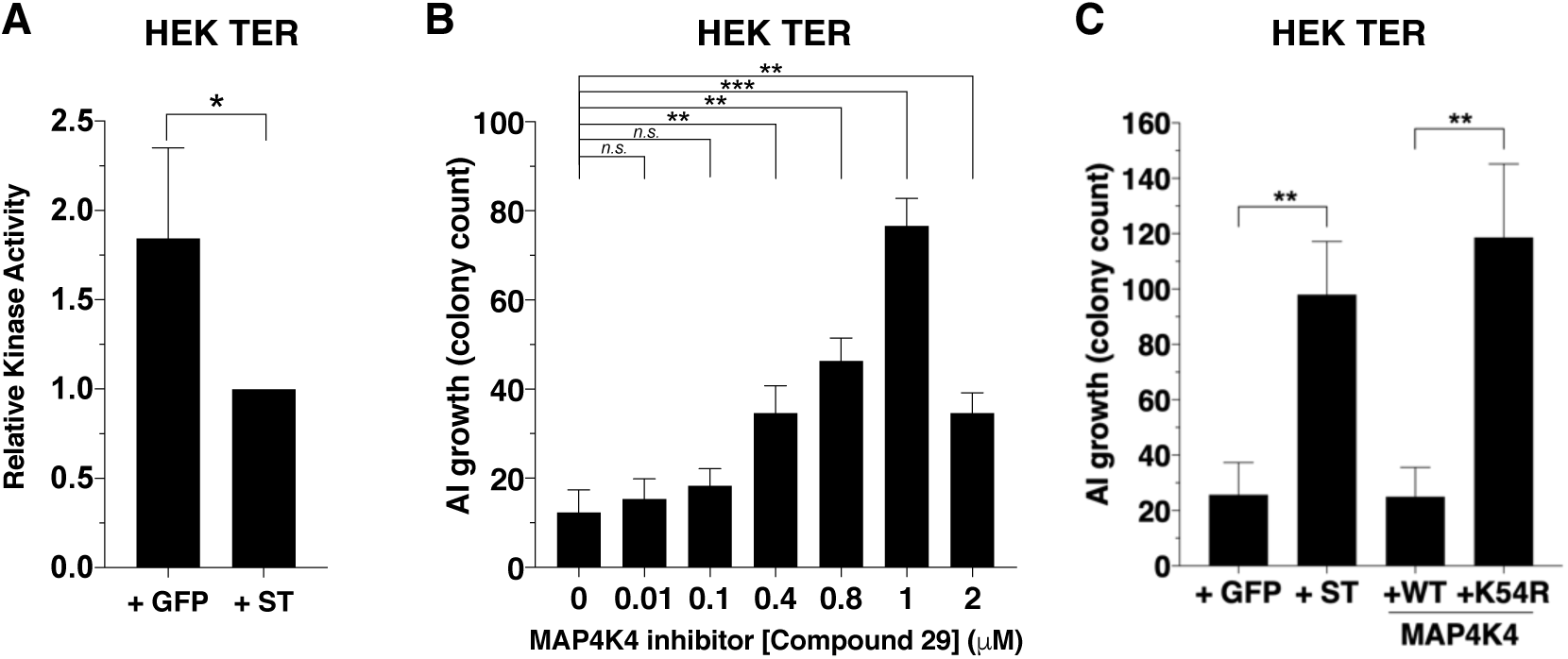
Partial inhibition of MAP4K4 kinase activity elicits transformation. (A) Quantification of MAP4K4 *in vitro* kinase activity after MAP4K4 was tandem-affinity purified from cells expressing ST or GFP control. (B) Quantification of AI growth after increasing concentrations of the MAP4K4 inhibitor C29. (C) Quantification of AI growth after expression of MAP4K4 WT, K54R, ST, or GFP in HEK TER cells (Student’s t-test, *p<0.01, **p<0.001, ***p<0.0001, n.s.= not significant).

To assess the relevance of MAP4K4 kinase activity to the transformation phenotype, we tested the consequences of pharmacological or genetic inhibition of MAP4K4 on AI growth. Specifically, we treated HEK TER cells with a previously described small molecule inhibitor of MAP4K4 (compound 29) (Crawford et al. 2014) over a range of concentrations (0-2μM) and assessed MAP4K4 activity (Fig. S4A) and AI cell growth (Fig. 4B). In consonance with what we observed with partially knocked down MAP4K4 expression, escalating doses of this MAP4K4 inhibitor led to an increase in the number of AI colonies until it reached 2 μM when MAP4K4 kinase activity was inhibited more than 90% as measured by *in vitro* kinase assays (2 μM, Fig. 4B, S4A). We found that compound 29 induced modest effects on cell proliferation over the range of tested concentrations (Fig. S4B). Consistent with the results from the genetic experiments (Fig 2B-C), we observed that partial inhibition of MAP4K4 activity led to increased AI growth.

We also tested whether inhibiting MAP4K4 by expressing a loss-of-function MAP4K4 allele promoted transformation. The kinase dead MAP4K4 K54R allele has previously been demonstrated to act as a dominant interfering mutant (Wang et al. 2013). We created HEK TER cells stably expressing kinase dead (K54R) or the wild-type (WT) version of MAP4K4 and confirmed the loss of kinase activity for the MAP4K4 mutant allele (Fig. S4C-D). When we performed AI growth assay, we observed that the introduction of MAP4K4 K54R but not WT MAP4K4 induced cell transformation (Fig. 4C). Together, these observations demonstrate that partial depletion or inhibition of MAP4K4 activity mimics ST in inducing transformation and that attenuation of MAP4K4 kinase activity is associated with ST-induced cell transformation.

### STRN4 is required for ST-mediated transformation

Reduction of MAP4K4 levels and activity was sufficient to drive transformation in the absence of ST; therefore, we also investigated whether members of the STRIPAK complex were required for ST-mediated oncogenic transformation (Fig. 5A). Specifically, we assessed the consequences of depleting components of STRIPAK in HER TER ST cell and found that knockdown of STRN4 led to significant reduction in transformation (Fig. 5B-C). We tested 4 STRN4-targeting shRNAs and observed reduction in AI colonies in a manner that significantly correlated with the degree of STRN4 knockdown (Fig. S5A-B). To confirm that these findings were not due to an off-target effect of RNAi, we created a STRN4 allele (STRN4-58R) resistant to the STRN4-specific shRNA (shSTRN4-58) and expressed this in HEK TER ST cells (Fig. S5C). We found that expression of this STRN4 allele rescued the effects of suppressing STRN4 on AI growth (Fig. 5D). We also deleted STRN4 using CRISPR-Cas9 gene editing and further confirmed that STRN4 expression was required for ST induced cell transformation (Fig. S5D). We assessed the consequences of knocking down STRN4 *in vivo*, and found that STRN4 knockdown significantly reduced tumor formation of HEK TER ST cells (Fig. 5E-F). Collectively, these observations demonstrate that STRN4 is required for ST-mediated transformation and tumor formation.

**Figure 5.**
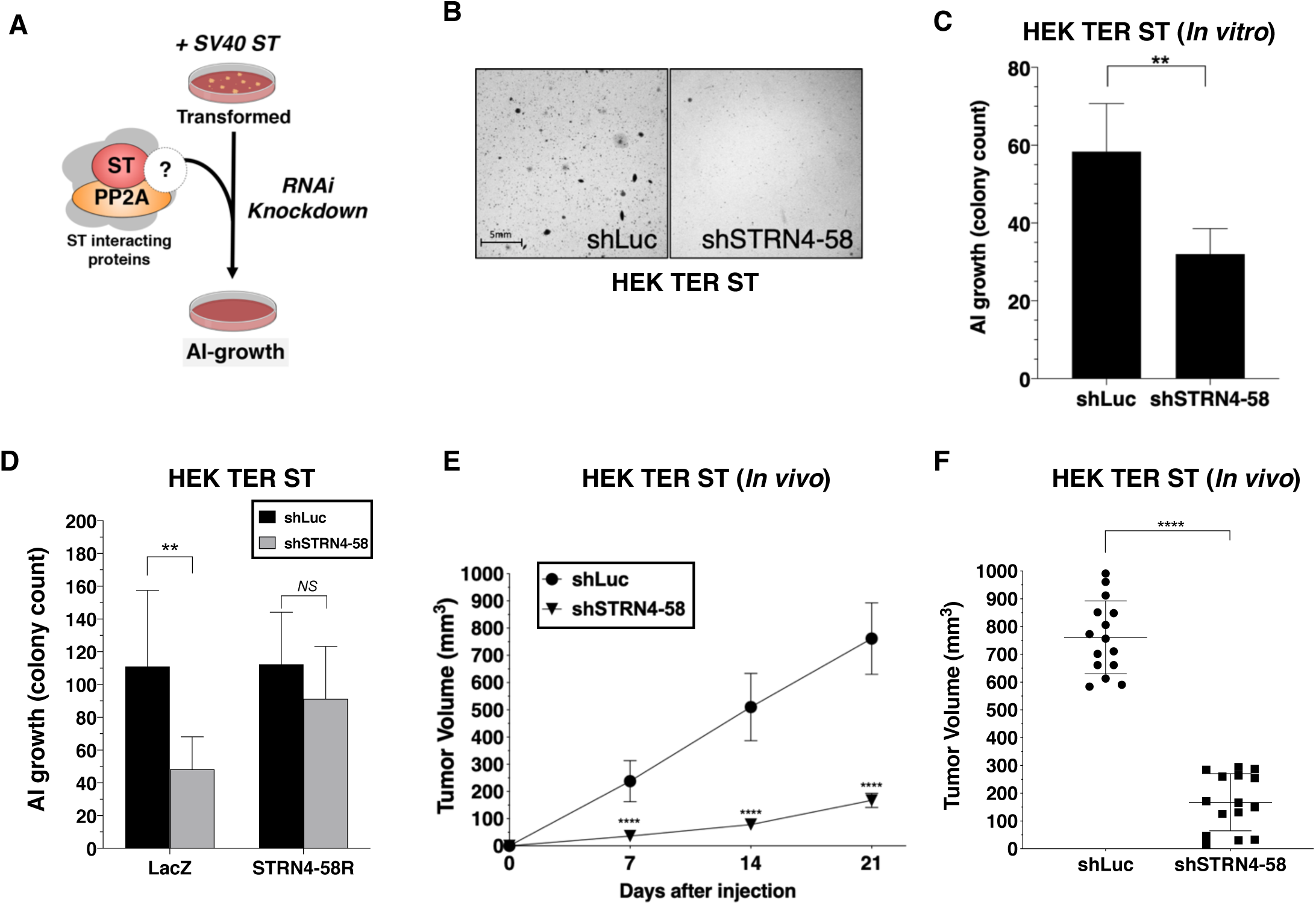
STRN4 is required for ST-mediated transformation and tumor induction. (A) Schematic of experimental design to reveal binding proteins that when depleted inhibit ST-mediated transformation. (B) Representative images of AI colonies observed after knockdown of STRN4 in HEK TER ST cells with shSTRN4-58. (C) Quantification of the number of AI colonies following introduction of shSTRN4-58 or shLuc control. (D) Quantification of the number of AI colonies after expression of shSTRN4-58 in the presence (STRN4-58R) or absence (LacZ) of an shRNA-resistant STRN4 cDNA. Tumor volume as a function of time (E) or at the endpoint at day 21 (F) for subcutaneous xenografts expressing shLuc control or STRN4 shRNA (shSTRN4-58) in HEK TER ST cells (Student’s t-test, **p<0.001, ****p<0.00001).

### STRN4 is required for the STRIPAK complex to associate with MAP4K4

To assess whether ST modulates interactions involving STRN4, we isolated endogenous STRN4 from cells expressing either ST or a GFP control and performed a proteomic analysis of associated proteins (Fig. 6A). We found that STRN4 interactions with MAP4K4 and with PP2A Cα were increased 2 and 3-fold, respectively, in cells expressing SV40 ST relative to GFP control (Fig. 6B).

**Figure 6.**
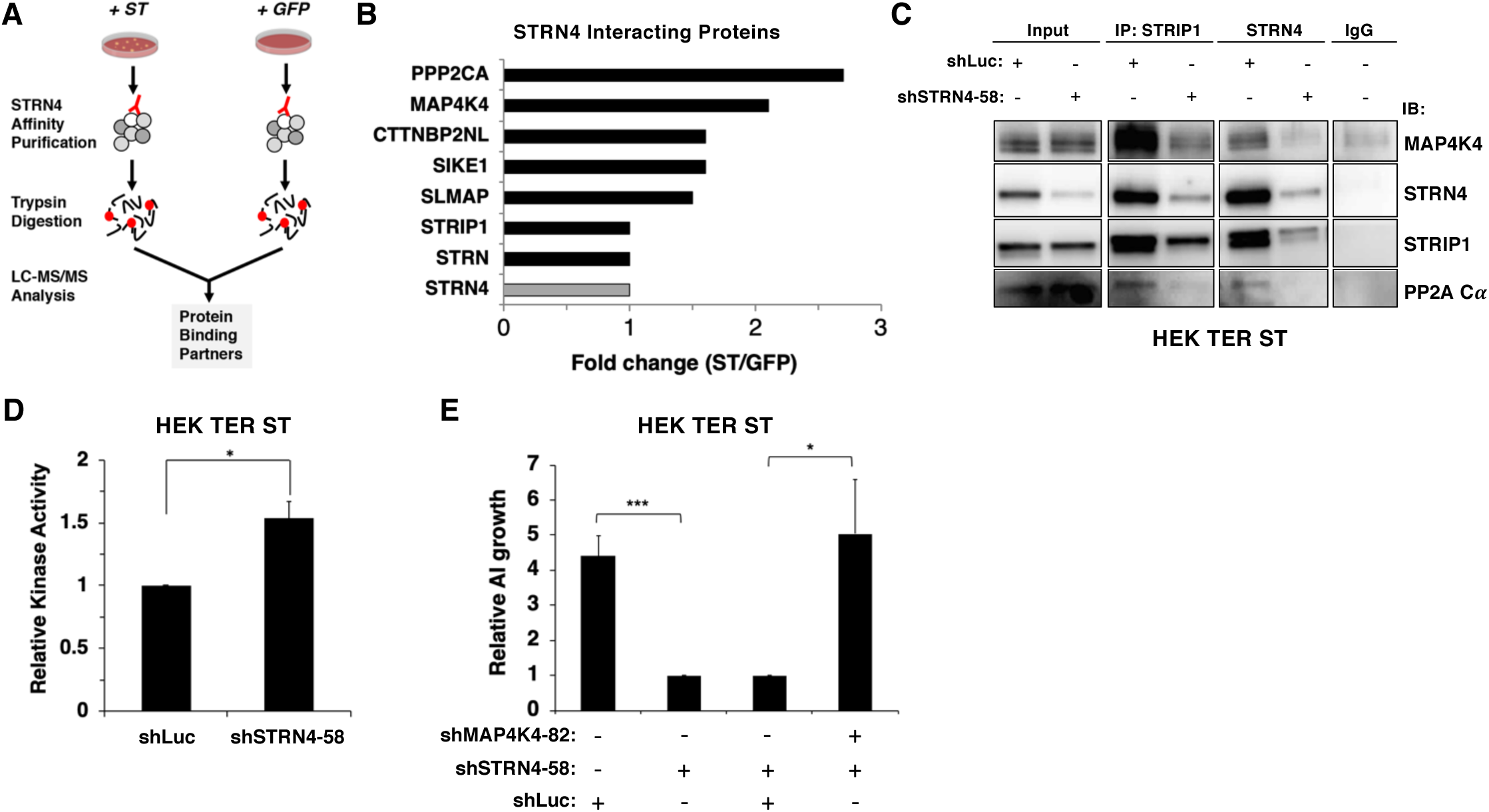
STRN4 is required for STRIPAK to interact with MAP4K4. (A) Schematic of proteomic analysis of STRN4 interacting proteins in the presence of ST or a GFP control. (B) Fold change in abundance of STRN4 interacting proteins in the presence of ST compared to the GFP control. All values were normalized to STRN4 levels to account for variations in the total amount of STRN4 isolated from cells expression ST or GFP control. Fold-change was calculated to reflect differences in the amount of STRN4 interacting proteins between cells expressing ST relative to GFP. Interactions of a number of proteins in the STRIPAK complex, including PP2A Cα, MAP4K4, CTTNBP2NL, SIKE1 and SLMAP with STRN4 were increased in ST-expressing cells relative to the GFP control. (C) Immunoblot showing a Co-IP analysis of STRIPAK core components and MAP4K4 after knockdown of STRN4 using shRNA. STRN4 is required for the STRIPAK component STRIP1 to interact with MAP4K4 and PP2A Cα. (D) Quantification of MAP4K4 *in vitro* kinase activity after STRN4 knockdown (shSTRN4-58). (E) Quantification of AI growth after expression of STRN4 shRNA with or without co-expression of MAP4K4 shRNA in HEK TER ST cells. Suppression of MAP4K4 expression rescued the transformation defect arising from STRN4 depletion. All experiments were performed in triplicate and the statistical analyses were performed relative to the controls (Student’s t-test, *p<0.01, ***p<0.0001).

Because ST promoted the interaction of MAP4K4 with STRN4, we evaluated the role of STRN4 in organizing the STRIPAK complex. When we assessed the impact of knocking down STRN4 on the STRIPAK complex in HEK TER ST cells by Co-IP of endogenous STRIP1, STRN4, and MAP4K4 (Fig. 6C, S6A) with or without STRN4 suppression, we observed that interactions of MAP4K4 with other members of STRIPAK (STRIP1, PP2A Cα) were attenuated when STRN4 was suppressed, indicating that STRN4 is required for MAP4K4 interactions with the STRIPAK complex.

Prior studies have shown that Striatins act as scaffolds in the STRIPAK complex (Chen et al. 2014). Based on these observations, we hypothesized that depletion of STRN4 in the presence of ST would lead to dissociation of MAP4K4 from the STRIPAK complex, which in turn would increase MAP4K4 activity. To test this hypothesis, we performed an *in vitro* kinase assay using MAP4K4 isolated from HEK TER ST cells expressing either control or STRN4-specific shRNA. We observed a modest, but statistically significant (p<0.05) increase in MAP4K4 kinase activity when STRN4 was suppressed (Fig. 6D, S6B). Moreover, we found that co-knockdown of MAP4K4 and STRN4, rescued the cells from the inhibitory effect of shSTRN4 knockdown on AI growth (Fig. 6E). In addition, we also observed that the expression of the dominant inhibitory K54R mutant, but not wild type MAP4K4, was able to restore the ability of these cells to form AI colonies upon STRN4 suppression (Fig. S6C). These observations suggest that ST inhibits MAP4K4 activity through STRN4 and the STRIPAK complex to induce transformation.

### Partial MAP4K4 knockdown induces YAP1 activation

To identify downstream signaling pathways affected during transformation by partial knockdown of MAP4K4 expression, we performed transcriptomic profiling of HEK TER cells expressing either shMAP4K4-82, which induced transformation in vitro (Fig. 2B-C), or a control shRNA targeting luciferase. We then performed single sample Gene Set Enrichment Analysis (ssGSEA) (Barbie et al. 2009) with the MAP4K4 knockdown gene expression signature and observed that several independent YAP1 genesets from the literature as well as two curated YAP1 and TAZ genesets from Ingenuity Pathway Analysis (IPA) were significantly associated with MAP4K4 knockdown (Fig. 7A) (Zhao et al. 2007, Yu et al. 2012, Hiemer et al. 2015, Martin et al. 2018). We also observed that phosphorylation of YAP1 at S127, a critical, negative regulatory site that blocks nuclear import of YAP1 (Zhao et al. 2007), was decreased upon partial knockdown of MAP4K4 or expression of the MAP4K4 K54R construct in HEK TER cells (Fig. 7B). Consistent with prior reports on regulation of LATS1/2 by MAP4K4 (Mohseni et al. 2014, Meng et al. 2015, Zheng et al. 2015), we also found that partial knockdown of MAP4K4 led to attenuation of p-LATS1 (Fig. 7C). In addition, we observed that mRNA and protein levels of CTGF and CYR61, established markers of YAP1 activity, were increased upon knockdown of MAP4K4 (Fig. 7D-F). These observations showed that partial knockdown of MAP4K4 at levels that induce cell transformation also led to increased YAP1 activity. In contrast, we found that suppression of STRN4 in HEK TER ST cells led to an increase in pYAP1 (Fig. 7G), consistent with the change in MAP4K4 activity upon STRN4 knockdown (Fig. 6D).

**Figure 7.**
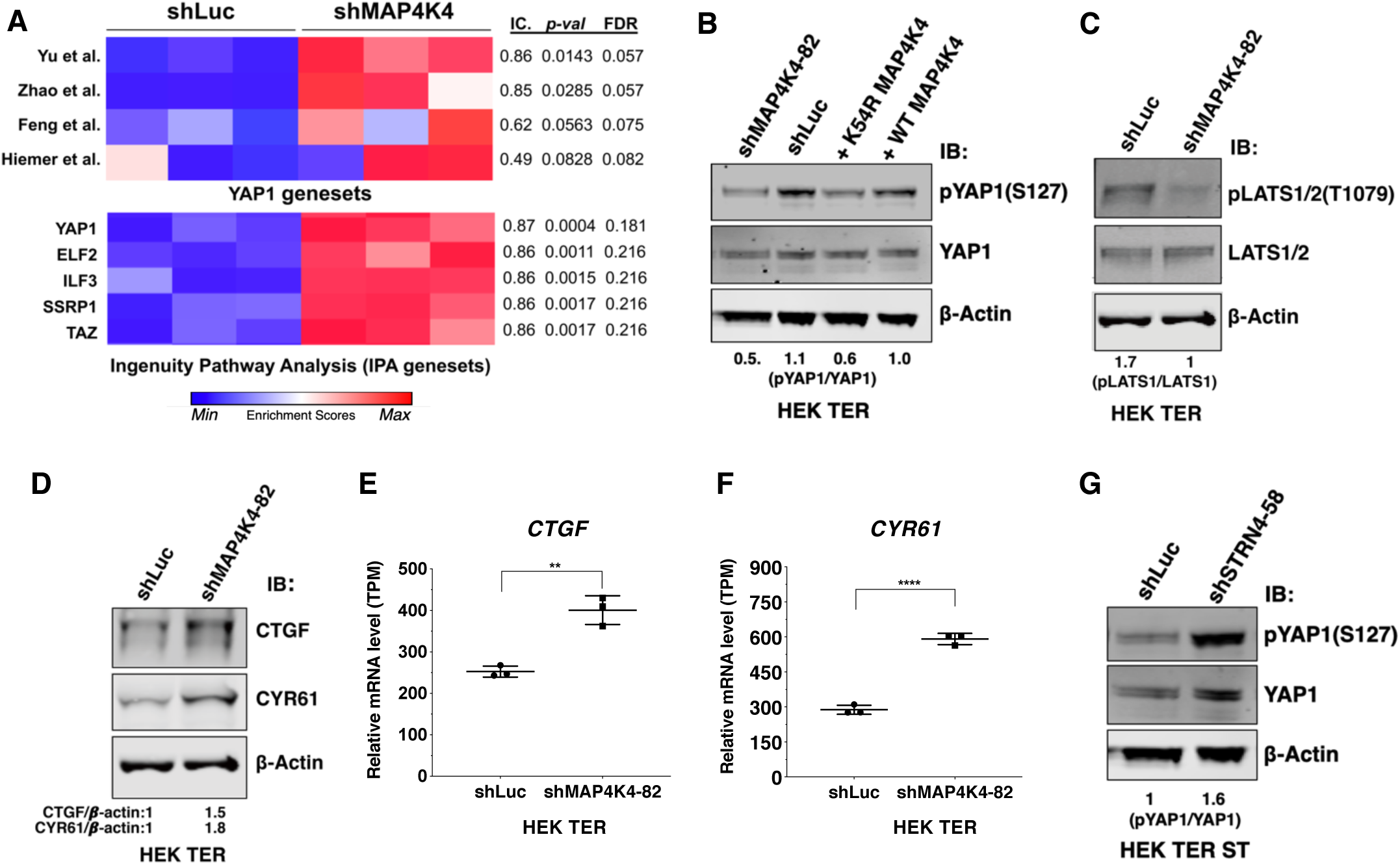
Depletion of MAP4K4 and STRN4 are linked to YAP1 regulation. (A) Heatmap of Enrichment Scores (ES) from RNA-seq analysis showing that partial suppression of MAP4K4 expression in HEK TER cells upregulates a transcriptional signature closely resembling four published, independently generated YAP1 signatures and two signatures for YAP1/TAZ from Ingenuity Pathway Analysis (IPA) using Information Coefficient (IC) as a similarity metric. ssGSEA was performed and enrichment scores are represented as indicated in the color bar with red indicating relative enrichment and blue depletion. The 3 columns in the heatmap represent triplicates for each condition. (B) Immunoblot depicting changes in phosphorylation of YAP1 on a key negative regulatory site (S127) following partial MAP4K4 knockdown or expression of MAP4K4 K54R in HEK TER cells. The values below the blot represent quantitation of the YAP1 pSer127 signal relative to total YAP1 from the immunoblot. (C) Immunoblot showing changes in phospo-LATS1 following partial MAP4K4 knockdown in HEK TER cells. Quantification of the LATS1 Thr1079 signal relative to total LATS1 from the immunoblot is shown below gel. (D) Immunoblot depicting changes in the YAP1 target genes CTGF and CYR61 following partial MAP4K4 knockdown and the ratios of the levels of CTGF/β-actin, CYR61/β-actin are shown below the blot. β-actin shown was performed in the same blot. Changes in the mRNA levels of YAP1 target genes CTGF (E) and CYR61 (F) upon MAP4K4 suppression. (G) Immunoblot depicting changes in phosphorylation of YAP1 on S127 following STRN4 knockdown in HEK TER ST. The values below the blot depict quantitation of the YAP1 pSer127 signal relative to total YAP1 from the blot (**p<0.001, ****p<0.00001).

### MAP4K4 activity converges on regulation of the Hippo/YAP1 pathway

To evaluate the role of YAP1 in transformation induced by attenuation of MAP4K4, we suppressed both MAP4K4 and YAP1 and tested AI colony formation (Fig. 8A). We found that although knockdown of MAP4K4 sufficed to promote transformation, we observed a three-fold decrease in AI colony growth when MAP4K4 was co-suppressed with YAP1 relative to an shRNA targeting luciferase as a control, indicating that transformation following partial knockdown of MAP4K4 depends on YAP1 (Fig. 8A, S7A).

**Figure 8.**
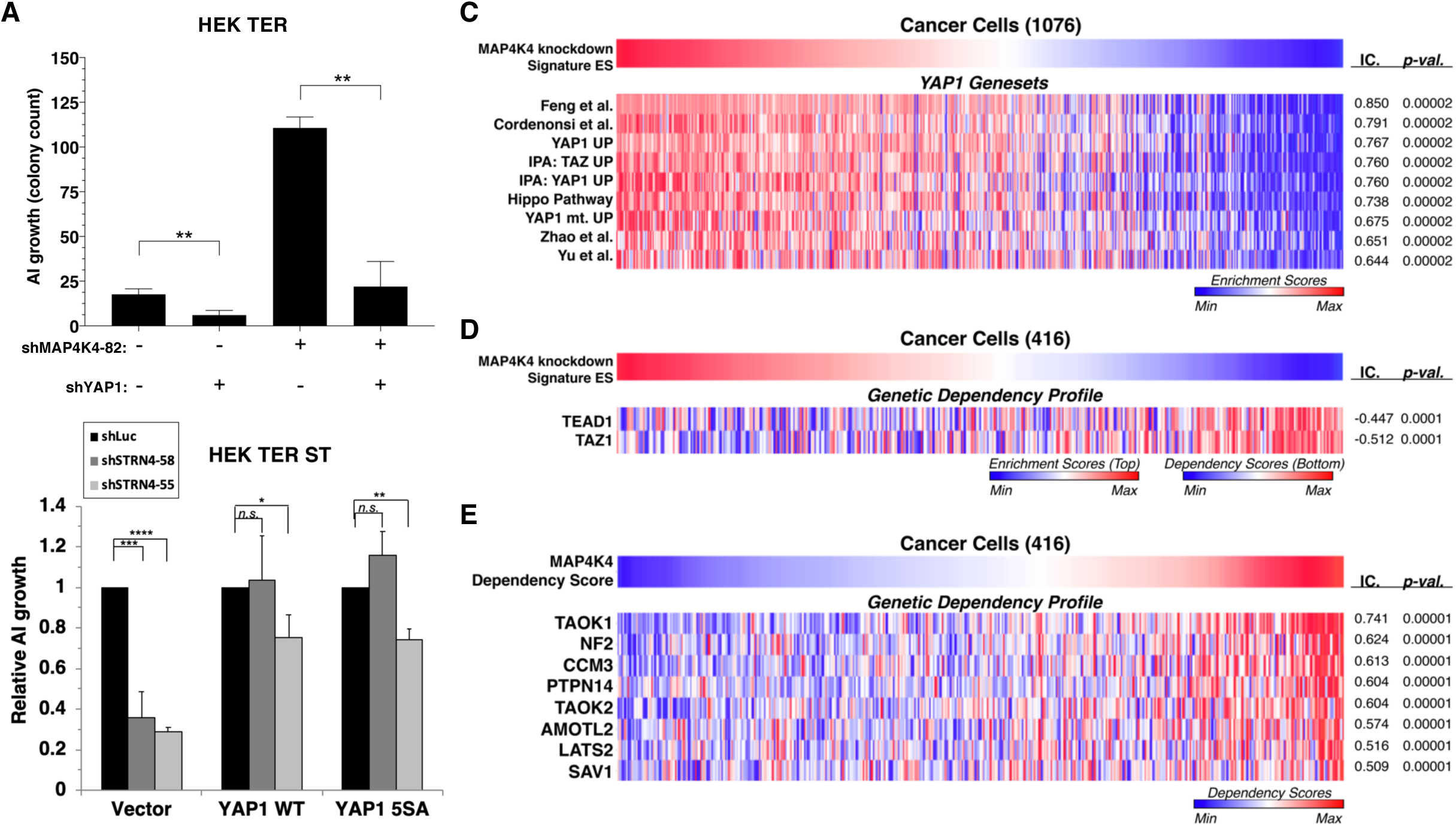
YAP1 is necessary for transformation upon MAP4K4 knockdown and rescues transformation in STRN4 knockdown cells. (A) Quantification of AI growth obtained following partial MAP4K4 suppression alone or when combined with YAP1 suppression (shYAP1) in HEK TER cells. Transformation induced by partial MAP4K4 suppression depends on YAP1. (B) Quantification of AI growth following *STRN4* knockdown with or without co-expression of YAP1 WT or the S5A mutant in HEK TER ST cells. YAP1 rescues the transformation defect of STRN4 suppression by shSTRN4-55 and shSTRN4-58 (immunoblots are shown in Fig. S7B). (C) Heatmap of ES depicting YAP1 genesets from the literature significantly associated with the MAP4K4 knockdown signature ES using Information Coefficient (IC) as a similarity metric. ssGSEA was performed using respective genesets across the CCLE dataset and enrichment scores are represented as indicated in the color bar, with red indicating relative enrichment and blue depletion (FDR<0.0001). (D) Heatmap depicting top dependency genes (bottom heatmap) in the Project Achilles dependency profiles that associated with MAP4K4 knockdown signature ES (top heatmap). Top heatmap represents ES from ssGSEA, while the bottom heatmap represent relative dependency with blue indicating strong dependency. Cell lines with low MAP4K4 transcriptional activity (in blue on top) were the most dependent on TEAD1 and TAZ1 (FDR<0.0001). (E) Heatmap depicting co-dependency analysis of MAP4K4 using IC across the Project Achilles data. The genes most significantly associated with MAP4K4 dependency were enriched for the Hippo/YAP1 pathway, as well as components of the STRIPAK complex as depicted (All associations FDR<0.0001 except TAOK2, SAV1: FDR=0.002). (Student’s t-test, *p<0.01, **p<0.001, ****p<0.00001, n.s.= not significant).

To further investigate the involvement of YAP1 activity in transformation, we tested whether expression of a constitutively active YAP1 phospho-mutant allele (5SA) (Zhao et al. 2007) rescued transformation when STRN4 was knocked down (Fig. S7B). We found that reduced levels of AI growth induced by STRN4 knockdown was rescued by expression of the wild-type or phospho-mutant YAP1 (Fig. 8B, S7B). These observations show that expression of YAP1 or YAP1 5SA overrides the requirement for STRN4 in transformation.

To extend these observations beyond the HEK TER cells, we generated a MAP4K4 knockdown gene expression signature and assessed this signature across a large collection of cancer cell lines from the Cancer Cell Line Encyclopedia (CCLE) by performing ssGSEA analysis (Fig. 8C-D) (Barbie et al. 2009, Barretina et al. 2012). Using the resulting Enrichment Scores (ES) derived from the MAP4K4 knockdown signature, we calculated information-theoretic measure, the Information Coefficient (IC) (Kim et al. 2016) to examine genesets that best matched the MAP4K4 knockdown signature ES across these cancer cell lines. In consonance with the findings in isogenic experiments (Fig. 7A), we observed that the MAP4K4 knockdown signature associated significantly with a number of YAP1 genesets derived from the literature as well as those we have generated by ectopic expression of wild-type or mutant YAP1 in immortalized human mammary epithelial cell (YAP1 UP, YAP1 mt UP) (Fig. 8C) (p-value<0.0001) (Zhao et al. 2008, Cordenonsi et al. 2011, Yu et al. 2012, Hiemer et al. 2015, Feng et al. 2019). Furthermore, when we compared the MAP4K4 knockdown signature with gene dependency data from Project Achilles, a large-scale project involving genome scale loss-of-function fitness screens performed in hundreds of cancer cell lines (Aguirre et al. 2016, Meyers et al. 2017, Tsherniak et al. 2017), we observed significant association with dependency profiles of TAZ1 and TEAD1 (IC= −0.447, −0.512, p-value= 0.0001, 0.0001, respectively), which are both Hippo pathway effector molecules (Fig. 8D). These findings indicated that the MAP4K4 knockdown signature associated with dependencies in the Hippo/YAP1 pathway.

We recently showed that systematically evaluating patterns of genetic co-dependencies across a dataset identify genes with similar function (Pan et al. 2018). We used this same approach to examine the MAP4K4 dependency profile. The MAP4K4 dependency profile quantitatively reflected the relative effect of targeting MAP4K4 on cell proliferation/survival of 416 cell lines and is represented as “dependency scores” on cell proliferation/survival during MAP4K4 inhibition (Fig. 8E). To assess genes that share dependency profiles with MAP4K4, we performed an orthogonal analysis using IC-based associations to identify a group of genes whose dependency profiles were most significantly associated with MAP4K4 dependency. Consistent with the known role of MAP4K4 in regulating the Hippo pathway, we found that a number of genes whose dependency profiles were most significantly associated with those of MAP4K4 belonged to the Hippo pathway, such as LATS2, PTPN14 and NF2 (Fig. 8E) (top 25 among the 18,375 dependency profiles)(p=0.00001). We also observed that CCM3 (PDCD10), a member of the STRIPAK complex (Goudreault et al. 2009), was the top most significantly associated gene dependency with MAP4K4, further supporting a link between MAP4K4, Hippo and the STRIPAK complex (Fig. 8E). These observations suggest that the gene expression associated with MAP4K4 knockdown is observed in many cancer cell lines and correlates with the Hippo signaling pathway.

### Discussion

Several lines of evidence now implicate the disruption of specific PP2A complexes and alteration of substrate specificity by mutation, deletion, or expression of polyomavirus ST as the basis for PP2A-mediated tumor suppressor activity. These observations have led to a model in which cancer associated PP2A mutations or ST alter the composition of PP2A complexes in cells, thus altering PP2A activity toward specific substrates. However, since purified PP2A exhibits phosphatase activity towards a broad set of substrates, the mechanisms that regulate PP2A substrate specificity in cells remains incompletely understood (Yang et al. 1991). Here we show that STRIPAK regulates the interaction of PP2A with one substrate MAP4K4 that participates in PP2A-dependent cell transformation. These observations provide a mechanism by which phosphatase activity is regulated.

Previous studies had shown that most B subunits were displaced by ST from the core enzyme and could not be detected in complex with ST (Pallas et al. 1990, Chen et al. 2007, Cho et al. 2007, Sablina et al. 2010). However, our proteomic analysis revealed that ST was bound to B’’’ subunits (striatins), as well as several other STRIPAK components. Here, we evaluated whether suppressing expression of these STRIPAK components impacted ST-induced cell transformation. We found that ST expression induced increased interactions of MAP4K4 with the STRIPAK complex, which in turn reduced levels of MAP4K4 phosphorylation and activity, thus leading to increased YAP1 activity. Prior studies have connected STRIPAK with components of the Hippo pathway (Couzens et al. 2013), and MAP4K4 has been shown to directly activate LATS1/2 kinases (Meng et al. 2015). In addition, YAP1 has been shown to be required for SV40 ST-mediated transformation (Nguyen et al. 2014). Recent work has shown mouse polyomavirus middle T affects YAP1 by directly binding to YAP1 and suppressing its degradation (Hwang et al. 2014, Rouleau et al. 2016). We propose a model for cell transformation induced by PP2A-mediated dephosphorylation of YAP1 in Fig. S7C.

We have shown that partial knockdown of MAP4K4 levels or inhibition of its kinase activity replaces ST in cell transformation, suggesting that MAP4K4 is a key PP2A substrate necessary for cell transformation. This observation is similar to our prior work that shows that only partial, but not complete, knockdown of PP2A Aα and Cα subunits leads to transformation (Chen et al. 2004, Chen et al. 2005, Sablina et al. 2010). However, we also note that the observed effects of suppressing MAP4K4 leads to greater increase in AI growth when compared to PP2A knockdown. We speculate that this may be due to the large repertoire of PP2A substrates that may have both pro-tumorigenic, as well as anti-tumorigenic activities. Likewise, MAP4K4 has been associated with a number of different pathways and biological processes (e.g. invasion, metabolism, TNF-α) and therefore, full depletion of MAP4K4 may impact other processes that are important for transformation. Although we observed increased interactions of MAP4K4 with the STRIPAK complex in cells expressing ST, only a subset of cellular MAP4K4 interacts with STRIPAK in this context (Fig. 3C), further supporting the notion that MAP4K4 unbound to the STRIPAK complex may have pro-tumorigenic roles. These observations also suggest that the STRIPAK complex plays a key role in regulating PP2A activity towards specific substrates and support a model in which ST in part induces transformation by promoting interactions of the STRIPAK complex with MAP4K4 and thereby attenuating MAP4K4 kinase activity, which in turn leads to the activation of YAP1.

The mechanism by which different PP2A complexes achieve substrate specificity has long remained elusive. Recent work has shown that proteins that harbor a conserved LxxIxE motif promote interactions with B56 subunits and facilitate subsequent PP2A substrate specificity (Hertz et al. 2016), suggesting that the substrate specificity may be achieved in part through specific interactions achieved by interactions with distinct B subunits. These findings reinforce the notion that STRIPAK serves as an organizing scaffold to bring substrates such as MAP4K4 to the PP2A complex. Indeed, recent studies have shown that MST3, a member of the STRIPAK complex, and Ste20 kinase family member MINK1 are also substrates of the STRIPAK complex (Gordon et al. 2011, Hyodo et al. 2012). Although we did not observe the presence of these proteins by proteomics in our current studies, it will be of interest to see if these proteins also affect transformation phenotypes in other contexts.

We found that the PP2A A-C complex continues to interact with non-canonical B’’’ subunits in the presence of ST. This observation confirms prior work that showed that both STRN and STRN3 binding do not overlap with canonical B subunit binding to Aα (Moreno et al. 2000). Furthermore, ST has been shown to be unable to compete with and displace B subunits from interacting with the PP2A core enzyme (Chen et al. 2007). Indeed, early observations involving biochemical characterization of the PP2A–ST complex showed that even in the absence of canonical B subunits, PP2A bound to ST dephosphorylated histone H1, suggesting that ST may alter substrate specificity of PP2A (Kamibayashi et al. 1994). Here we provide further evidence that ST alters substrate specificity by promoting MAP4K4 interaction with the STRIPAK complex. It is unclear if ST binding to PP2A Aα promotes active conformational changes that increase PP2A AC subunit affinity for STRN4, or if there is competition among the canonical and non-canonical B subunits to engage the PP2A core enzyme complex. However, it appears that ST interactions with STRIPAK are dependent on Aα, as ST mutants that failed to bind to Aα were also unable to bind to STRIPAK (Fig. S3A). In addition, it was recently shown that disruption of interactions between PP2A core enzyme and canonical B subunits by mutations in PP2A Aα (P179R, R18G) promotes PP2A interactions with members of the STRIPAK complex (Haesen et al. 2016), reinforcing the notion that ST phenocopies the effect of cancer-associated PP2A mutations. More generally, these observations suggest that striatins act as key regulators of PP2A that impart substrate specificity.

MAP4K4 is less well characterized than other members of the MAPK family but has been implicated in a number of biological processes including invasion, insulin resistance and immunity (Collins et al. 2006, Tang et al. 2006, Huang et al. 2014, Danai et al. 2015, Vitorino et al. 2015). Indeed, MAP4K4 has been reported to promote invasion and to act as a downstream component of TNF-α signaling (Wright et al. 2003, Crawford et al. 2014, Gao et al. 2016). However, others have found evidence that MAP4K4 can also act as a candidate tumor suppressor gene (Westbrook et al. 2005), promote apoptosis downstream of Sox2 (Chen et al. 2014, Yang et al. 2015) and serve as a regulator of the Hippo pathway, in part through direct phosphorylation of LATS1/2, leading to YAP/TAZ inhibition (Couzens et al. 2013, Mohseni et al. 2014, Meng et al. 2015, Zheng et al. 2015).

YAP1 is a downstream effector of the Hippo pathway and is involved in a number of important cellular processes including organ size control and cell proliferation. When the Hippo pathway is activated by upstream stimuli triggered by cell-cell contact, cell density and detachment, YAP1 is negatively regulated through a cascade of phosphorylation events causing YAP1 to reside in the cytoplasm and remain inactive. Therefore, tight regulation of the phosphorylation and dephosphorylation events that control the Hippo pathway and subsequent YAP1 activity is critical for preserving normal cellular homeostasis. YAP1 has also been shown to play prominent roles in oncogenic transformation, drug resistance and the epithelial-mesenchymal transition (Hong et al. 2014, Shao et al. 2014, Wilson et al. 2015). YAP1 has also been shown to be required for KRAS and ST-mediated transformation, providing further evidence that YAP1 is critical for cancer development and maintenance (Hong et al. 2014, Nguyen et al. 2014, Shao et al. 2014).

Despite clear evidence for YAP1 in both cancer initiation and progression, few mutations involving YAP1 or other Hippo pathway components have been identified in cancers. Since mutations affecting PP2A subunits are commonly observed in several types of cancer, our observation that certain PP2A complexes can activate YAP1 in the context of ST-mediated transformation suggests that these cancer-associated mutations may also serve, in part, to activate YAP1.

## METHODS

### Cell lines

HEK TER cells were generated from human embryonic kidney (HEK) cells, which were immortalized by introducing hTERT, SV40 Large-T antigen, and H-RAS G12V (Hahn et al. 2002). These cells were cultured in MEM-alpha media (Gibco) supplemented with 10% FBS. 293T cells (ATCC) and HCT-116 (ATCC) cells were cultured in Dulbecco’s modified Eagle medium (DMEM) (Cellgro) supplemented with 1% Pen-Strep (Gibco), 1% Glutamax (Gibco) and 10% fetal bovine serum (FBS) (Sigma). IMR90 cells (ATCC) were cultured in DMEM supplemented with 1% Pen-Strep, 1% Glutamax, and 1% non-essential amino acids (Gibco) and 15% FBS.

### Sample Preparations for the Global Phosphoproteomics

HEK TER cells expressing SV40ST or suppressed the expression of PP2A Cα, Aα or B56γ subunits were synchronized in serum-free medium for 24 h, followed by serum stimulation (5 min.) and immediately harvested. Experiments were performed on two independent days as replicates.

### Global Phosphoproteomics

Cell pellets were solubilized by repeated pipetting using in 10 volumes of 7.2M guanidine HCl 0.1M ammonium bicarbonate. Insoluble material was pelleted for 10 minutes at 10.000 x g and the protein concentration of the supernatants quantified by bicinchoninic acid assay (Pierce). Aliquots corresponding to 50 micrograms of each sample were transferred to new tubes and the volumes brought to 50 microliters using the above solubilization buffer before further processing. Cysteine residues were reduced with 10 mM dithiothreitol (DTT) for 30 minutes at 56°C and alkylated with 22.5 mM iodoacetamide for 20 minutes at room temperature in the dark. The concentration of guanidine HCl was lowered by adding 9 volumes of 0.1M ammonium bicarbonate. Samples were digested overnight at 37°C using 10 micrograms of trypsin (Promega). An additional 10 micrograms of trypsin were added the following morning and incubated for another 4 hours at 37°C. The resulting tryptic peptide solutions were acidified by adding trifluoroacetic acid (TFA) to a final concentration of 1% and desalted on a Waters C18 solid phase extraction plate (using two consecutive passes). Eluted peptides were concentrated in a vacuum concentrator and reconstituted with 30 µL of 0.5 M triethylammonium bicarbonate. Each tube of iTRAQ reagent was reconstituted with 70 µL ethanol and added to each peptide solution. The labeling reaction was carried out for 1 hour at room temperature. Labeled peptides were combined in a tube containing 100 µL of 16.5 M acetic acid, concentrated by vacuum centrifugation and desalted on a Waters C18 solid phase extraction plate. Magnetic Fe-NTA agarose beads (300 µL of a 5% bead suspension) were prepared as described (Ficarro et al. 2009). The beads were added to iTRAQ labeled peptides reconstituted with 80% acetonitrile / 0.1% TFA at a concentration of 0.5 µg/µL. enriched for 30 minutes at room temperature with end-over-end rotation. After removing the supernatant, beads were washed three times with 400 µL 80% acetonitrile / 0.1% TFA, and once with 400 µL of 0.01% acetic acid. Phosphopeptides were eluted for 5 minutes at room temperature with 50 µL of 0.75M ammonium hydroxide containing 100 mM EDTA. The beads were washed once with 50 µL of water and this wash was combined with the eluate. Eluted phosphopeptides were concentrated to 10 µL by vacuum centrifugation. Ammonium formate (pH10) was added to yield a final concentration of 20 mM. Enriched phosphopeptides were analyzed by multidimensional RP-SAX-RP-MS/MS (Ficarro et al. 2009) at a depth of 43 fractions on an LTQ-velos mass spectrometer. The spectrometer was operated in data dependent mode where the top 10 most abundant ions in each MS scan were subjected to alternating CAD (electron multiplier detection, 35% normalized collision energy, q=0.25) and HCD (image current detection, 45% normalized collision energy) MS/MS scans (isolation width = 2.0 Da (CAD) and 2.4 Da (HCD), threshold = 20,000). Dynamic exclusion was enabled with a repeat count of 1 and exclusion duration of 30 seconds. ESI voltage was 2.2 kV. MS spectra were recalibrated using the background ion (Si(CH3)2O)6 at m/z 445.12 +/- 0.03 and converted into a Mascot generic file format (.mgf) using multiplierz scripts (Askenazi et al. 2009, Parikh et al. 2009). CAD and HCD spectra were independently searched using both Mascot (version 2.3) and Protein Pilot (version 4.5) against three appended databases consisting of: i) human protein sequences (downloaded from RefSeq on 07/11/2011); ii) common lab contaminants and iv) a decoy database generated by reversing the sequences from these two databases. For Mascot searches, precursor tolerance was set to 1 Da and product ion tolerance to 0.6 Da (CAD) or 0.02 Da (HCD). We used the default settings specified for CAD and HCD spectra for Protein Pilot searches (with no precursor tolerance specified). Mascot search parameters included trypsin specificity, up to 2 missed cleavages, fixed carbamidomethylation (C, +57 Da) and iTRAQ8plex derivatization (K and N-terminus), variable oxidation (M, +16 Da) and phosphorylation (S, T, Y, +80 Da). Protein Pilot search parameters included trypsin specificity, fixed carbamidomethylation (C, +57 Da), peptide level iTRAQ8plex labeling. mgf files corresponding to the 43 RP-SAX-RP MS/MS fractions were individually searched with Mascot and combined into one Excel file before calculating false discovery rate (FDR). Peptide summaries were exported as text files from the Protein Pilot search results and imported into Excel for FDR calculation. Data files were processed to remove i) peptide spectral matches (PSMs) to the reverse database; PSMs to the forward database with an FDR greater than 1.0% and iii) PSMs corresponding to spectrum with no iTRAQ reporter ions. PSMs were then compared across the 4 search results (Mascot CAD, Mascot HCD, Pilot CAD and Pilot HCD). PSMs with discordant peptide sequences were discarded. Peptide-level phosphorylation sites were selected based on a majority rule across searches for which an ID was made and used to locate protein-level phosphorylation in the SwissProt database (downloaded 02/06/2013; note that the entry for MAP4K4 (O95819-2) in this database contained a deletion at S627). iTRAQ intensities were summed across all PSMs with peptide sequences overlapping the protein-level phosphorylation site. The screen was performed across two replicates after randomizing the assignment of iTRAQ channel to biological samples.

### Virus Production

Packaging and envelope plasmids were co-transfected with lentiviral or retroviral expression vectors into 293T cells using Lipofectamine 2000 (Life Technologies). Two days after transfection, 293T cell supernatant was clarified with a 0.45 μm filter and supplemented with 4 μg/mL polybrene (Santa Cruz) before transducing recipient cells. Stable cell lines were generated after selection with 2 μg/mL puromycin (Sigma), 5 μg/mL blasticidin (Invivogen), 500 μg/mL G418 (Sigma) and 50 μg/mL hygromycin (Santa Cruz) as required by each vector. For MAP4K4 inhibitor experiments, dimethyl sulfoxide (DMSO) (Sigma) or inhibitor (compound 29) (Crawford et al. 2014) was used at the indicated concentrations.

### Recombinant DNA constructs

MAP4K4 cDNA was generated by PCR-based Gateway cloning (Invitrogen) from HEK TER cells. NTAP-SV40 ST and NTAP-GFP have been previously described (Rozenblatt-Rosen et al. 2012). Mutations in MAP4K4 and SV40 ST were introduced using the QuikChange XL II site-directed mutagenesis kit (Agilent). Lentiviral shRNA constructs were obtained from the Genetic Perturbation Platform (GPP) at the Broad Institute (Cambridge, MA) (http://www.broadinstitute.org/rnai/public/). The following clone IDs were used for STRN3: TRCN0000365162, TRCN0000370206, STRIP1; TRCN0000164502, TRCN0000162951, MARCKS: TRCN0000197145, TRCN0000029041, and STK24: TRCN0000000641, TRCN0000000644. STRN4: TRCN0000036954 (shSTRN4-54), TRCN0000036955 (shSTRN4-55), TRCN0000036957 (shSTRN4-57), TRCN0000036958 (shSTRN4-58). Subsequent functional studies were carried out with TRCN0000036958 (shSTRN4-58) and TRCN0000036955 (shSTRN4-55) and as these were further confirmed to have stronger knockdown of STRN4 protein, as well as relatively less off-target effects. STRN4 open-reading frame (ORF) construct which is resistant to STRN4 shRNA (TRCN0000036958 or shSTRN4-58) was generated by cloning of the following sequence; GCCCTTGAAGTCGAACCAATTCATGCT, which was obtained from IDT as gblocks gene fragment (Integrated DNA Technologies), into STRN4 wild-type ORF in pdonr223 using Gibson assembly cloning kit (Cat#2611, New England Biolabs), followed by gateway cloning into pLX304 vectors from the GPP. For knockdown of MAP4K4, we used clone IDs: TRCN0000220092, TRCN0000220093 and TRCN0000195258, TRCN0000219681, TRCN0000219682, TRCN0000195121, TRCN0000199325. For most of the study, we focused on TRCN0000219682 (shMAP4K4-82) unless otherwise indicated, as described in the main text. For YAP1 shRNA, we used clone ID: TRCN0000107265.

For global phosphoproteomic experiments, we used PP2A shRNAs as previously described (Sablina et al. 2007). Specifically, shRNAs targeting PP2A Cα, Aα, B56γ subunits were obtained from Genetic Perturbation Platform (GPP) with the clone IDs: TRCN0000002483 (shPP2A Cα1), TRCN0000002484 (shPP2A Cα2), TRCN0000002494 (shPP2A B56γ1), TRCN0000002496 (shPP2A B56γ2) and TRCN0000231508 (shPP2A Aα). For STRN4 CRISPR-CAS9-mediated knockout, the lentiCRISPRv2 vector was used [a gift from Feng Zhang (Addgene plasmid # 52961) (Sanjana et al. 2014). STRN4-specific sgRNA sequences were obtained from the Avana library (Doench et al. 2016) and sgRNAs were cloned according to Zhang lab protocols (http://genome-engineering.org/gecko/wp-content/uploads/2013/12/lentiCRISPRv2-and-lentiGuide-oligo-cloning-protocol.pdf).

Gateway-compatible cDNA entry clones were transferred from pDONR221 or pDONR223 donor vectors to the respective retro- or lentiviral Gateway destination vectors via Gateway recombinational cloning (Life Technologies). The vectors MSCV-N-terminal-Flag-HA-IRES-PURO (NTAP) and MSCV-C-terminal-Flag-HA-IRES-PURO (CTAP), as well as all HPyV cDNAs, have been previously described (Sowa et al. 2009, Rozenblatt-Rosen et al. 2012, Berrios et al. 2015). Where indicated, untagged constructs were expressed in the CTAP vector with a TAA stop codon to exclude expression of the epitope tag. Wild-type or phospho-mutant YAP1 was cloned into pMSCV puro vector (Clontech) to generate pMSCV puro YAP1 WT or 5SA.

The following plasmids were obtained from Addgene: pBabe-hygro-hTERT (plasmid # 1773) (Counter et al. 1998), pBabe-HcRed-Ras (plasmid # 10678) (Boehm et al. 2005), pBabe-neo-large T cDNA (plasmid # 1780) (Hahn et al. 2002), pWZL-Blast-ST (plasmid # 13805) (Chen et al. 2004), lentiviral packaging plasmid psPAX2 and envelope plasmid pMD2.G (plasmid #12260, #12259), retroviral packaging plasmid pUMVC3 (plasmid # 8449) (Stewart et al. 2003), and envelope plasmid pHCMV-AmphoEnv (plasmid # 15799) (Sena-Esteves et al. 2004).

### Proteomic analysis of immunopurified MAP4K4

FLAG-MAP4K4 immuno-precipitates (Adelmant et al. 2019) were diluted in 100 mM Ammonium Bicarbonate containing 0.1% RapiGest (final concentration) and reduced with 10 mM DTT for 30 minutes at 56°C. Reduced cysteine residues were alkylated with 22.5 mM iodoacetamide for 20 minutes in the dark. Proteins were digested with 5 µg of trypsin overnight at 37°C. Tryptic peptides were purified by batch-mode reverse-phase chromatography (POROS 50R2, Applied Biosystems) and subjected to immobilized metal affinity chromatography (IMAC) to enrich phosphopeptides as described for the global phosphoproteomics screen. Peptides from the IMAC supernatant were concentrated under vacuum and purified by batch-mode strong cation exchange chromatography (POROS 50HS). Phosphopeptides were analyzed by LC-MS/MS as follow: Phosphopeptides were loaded off-line onto a precolumn (4 cm POROS 10R2) and eluted with an HPLC gradient (NanoAcquity UPLC system, Waters; 5%–40% B in 45 min; A = 0.2 M acetic acid in water, B = 0.2 M acetic acid in acetonitrile). Peptides were resolved on a self-packed analytical column (50 cm Monitor C18, Column Engineering) and introduced in the mass spectrometer (QExactive HF mass spectrometer, Thermo) equipped with a Digital PicoView electrospray source platform (New Objective). The mass spectrometer was operated in data dependent mode where the top 10 most abundant ions in each MS scan were subjected to high energy collision induced dissociation (HCD, 27% normalized collision energy) and subjected to MS/MS scans (isolation width = 1.5 Da, intensity threshold = 25.000, MS1 resolution: 120 000). Dynamic exclusion was enabled with an exclusion duration of 30 seconds. ESI voltage was set to 3.8 kV. Dedicated MS/MS scans were also included to continuously monitor precursors for two phosphopeptides identified in a previous analysis. Peptides from the supernatant were separated using a 90-minute HPLC gradient and analyzed using in the mass spectrometer as described above. MS spectra were recalibrated using the background ion (Si(CH3)2O)6 at m/z 445.12 +/- 0.03 and converted into a Mascot generic file format (.mgf) using multiplierz scripts. Spectra were searched using Mascot (version 2.6) against three appended databases consisting of: i) human protein sequences (downloaded from RefSeq on 06/26/2019); ii) common lab contaminants and iii) a decoy database generated by reversing the sequences from these two databases. Precursor tolerance was set to 20 ppm and product ion tolerance was set to 25 mmu. Search parameters included trypsin specificity, up to 2 missed cleavages, fixed carbamidomethylation (C, +57 Da) and variable oxidation (M, +16 Da) and phosphorylation (S, T, +80 Da). Spectra matching to peptides from the reverse database were used to calculate a global false discovery rate and were discarded. The intensity of heavy and light SILAC features was directly retrieved from the mass spectrometry raw files using the multiplierz python environment (Alexander et al. 2017). MAP4K4 phosphorylation sites were remapped to isoform 6 (UniProt accession O95819-6) and SILAC intensities were summed for individual sites identified across overlapping peptide sequences. The SILAC intensity ratio representing the relative abundance of phosphorylation sites in cells expressing GFP or ST was normalized to correct for small difference in immunopurified MAP4K4 in the respective samples as measured in the IMAC supernatant. The relative abundance of proteins in the IMAC supernatant was calculated by summing the intensities of the heavy or light features across peptides mapping uniquely to a gene (Askenazi et al. 2010). Two independent FLAG-MAP4K4 immunoprecipitations were performed on combined extracts of GFP and ST expressing cells metabolically encoded with heavy and light or light and heavy SILAC labels, respectively.

### Immunoprecipitation and immunoblotting

Cell lysates were obtained using lysis buffer (150 mM NaCl, 50 mM Tris-HCl, 1 mM EDTA, 0.5% NP-40, 10% glycerol, and protease and phosphatase inhibitor cocktail sets (Calbiochem)). Immunoprecipitations were performed with protein A/G magnetic beads (Millipore) mixed with immunoprecipitation antibodies. After overnight incubation at 4°C, beads were washed with high salt lysis buffer (containing 300 mM NaCl), boiled in SDS sample buffer (Boston BioProducts), resolved by SDS-PAGE (Criterion TGX precast gels, Bio-Rad), transferred to nitrocellulose membranes (Bio-Rad), blocked and incubated with the appropriate primary antibody in TBS-T overnight at 4°C. Detection of proteins was performed with horseradish-peroxidase conjugated secondary antibodies (Rockland), developed using Clarity Western ECL substrate (Bio-Rad), and imaged with a G:BOX Chemi detection system (Syngene).

### MudPIT

HEK TER cells expressing either SV40 ST or GFP (30 x 15-cm diameter plates) were harvested with lysis buffer and clarified cell extract was incubated overnight at 4°C with 20-100 μg antibodies crosslinked to 30 mg protein A agarose beads (Thermo Scientific) by dimethyl pimelimidate (DMP). Beads were washed with high salt lysis buffer 5 times, washed with TBS 2 times, and then eluted with 0.2 M glycine pH 3 and neutralized with 1 M Tris-HCl pH 8.0. Proteins were precipitated with trichloroacetic acid (20% final concentration) overnight at 4°C, washed with cold acetone and processed for subsequent MudPIT analysis, as previously described (Florens et al. 2006).

### *In vitro* kinase assay

TAP-purified MAP4K4 eluted in standard lysis buffer with protease and phosphatase inhibitors was added to kinase assay buffer (25 mM Tris-HCl pH 7.5, 5 mM β-glycerophosphate, 2 mM dithiothreitol, 0.1 mM sodium orthovanadate and 10 mM MgCl_2_) containing 20 μM ATP*γ*S (Abcam) and 1 μg of myelin basic protein (MBP) (Sigma). Where specified, ATP*γ*S was left out of the reaction as a negative control. Kinase reactions were carried out as previously described (Allen et al. 2007). Reactions were carried out at 30°C for 30 min. P-nitrobenzyl mesylate (PNBM) (Abcam) was then added (2.5 mM final) and the reaction was incubated at room temperature for 2 hours, followed by addition of 6x SDS loading buffer, boiling of samples, SDS-PAGE and subsequent immunoblotting for phosphorylated MBP. Relative activity was calculated as the ratio of the band intensities (measured with ImageJ) between the thiophosphate ester signal (phospho-MBP) and HA signal (NTAP-MAP4K4).

### PP2A phosphatase assay

To measure PP2A phosphatase activity, we used a PP2A Immunoprecipitation Phosphatase Assay Kit (Millipore Sigma, catalog number 17-313). In brief, HEK TER GFP or ST cells were lysed in 20 mM imidazole HCl, 2 mM EDTA, 2 mM EGTA, pH 7.0 with 10 μg/mL each of aprotinin, leupeptin, pepstatin, 1 mM benzamidine, and 1 mM PMSF. Two milligrams of the lysates were then immunoprecipitated with 2 μg of anti-STRN4 antibody (Abcam, ab177155) and 40 μl of protein-A-agarose beads at 4°C overnight. Beads were washed three times with lysis buffer followed by the Ser/Thr assay buffer. Phosphatase reactions were then performed in Ser/Thr assay buffer with a final concentration of 750 μM of MAP4K4 phosphopeptides: S771/ S775 (A-A-S-pS-L-N-L-pS-N-G-E-T-E-S-V-K), S876 (L-T-A-N-E-T-Q-pS-A-S-S-T-L-Q-K) or S1251 (V-F-F-A-pS-V-R-S) for 10 minutes at 30°C. To provide evidence that the immunoprecipitated phosphatase activity is PP2A, we treated parallel immunoprecipitates with 5 nM of okadaic acid (Cell Signaling, #5934). Dephosphorylation of the phosphopeptide was measured through malachite green phosphate detection at 650 nm.

### AI growth and proliferation assays

HEK TER AI growth in soft agar was performed as previously described (Hahn et al. 2002) using 6-well dishes with BactoAgar (Gibco) at concentrations of 0.3% top and 0.6% bottom layers. Wells were fed with top agarose once per week. After 4 to 5 weeks, cells were stained with 0.005% crystal violet (Sigma) in PBS and colonies were counted. For MAP4K4 inhibitor experiments, dimethyl sulfoxide (DMSO) (Sigma) or inhibitor (compound 29) (Crawford et al. 2014) were used at the indicated concentrations in both the bottom and top soft agar layers and included in refeedings. For proliferation assays, cells were seeded in triplicate in 24-well plates (day 0; 5 x 10^3^ cells per well). Cell density was measured by crystal violet assay at intervals after plating as previously described (Rozenblatt-Rosen et al. 2012).

### *In Vivo* Xenografts

For in vivo xenograft experiments, 2 x 10^6^ HEK TER (expressing shLuc or shMAP4K4-82) or HEK TER ST (expressing shLuc or shSTRN4-58) cells were subcutaneously injected into the top, left and right flanks of 5 female Taconic NCR-nude (CrTac:NCr-Foxn1nu) mice. For the shSTRN4 experiments, we re-engineered the HEK TER cells to express KRAS G12V, because other vectors containing HRAS G12V with various selection markers failed to produce sufficient levels of expression. Tumor volume was assessed via caliper measurement every week by the formula: volume = length x width^2^ x 0.5. All procedures were performed according to protocols approved by the Institutional Animal Care and Use Committees of the Dana-Farber Cancer Institute.

### RNA-sequencing

A total of 500,000 cells of either HEK TER shLuc or shMAP4K4-82 were seeded in three 15 cm dishes and allowed to grow for 48 hours. Total RNA was extracted using a RNeasy Plus Kit (Qiagen). RNA sequencing libraries were prepared using a NEBNext Ultra Directional RNA Library Prep Kit for Illumina, NEB E7420. The concentration of each cDNA library was quantified with the KAPA Illumina ABI Quantification Kit (Kapa Biosystems). Libraries were pooled for sequencing using the HiSeq 2500.

### Data Analysis

Global phosphoproteomic data: Data from iTRAQ experiments were processed by first merging the two replicate datasets, which resulted in 6025 phosphopeptides corresponding to 2465 individual proteins. We then normalized the raw read counts of each sample to the corresponding control experiments (shRNA against luciferase for shRNA experiments, and GFP for ST experiments) followed by log_2_ transformation. The resulting values were further normalized by quantile normalization. We performed comparative marker selection to find phosphorylation changes which are most significantly correlated with cell transformation phenotype using signal-to-ratio statistics after 1000 permutations (Gould et al. 2006). The transformation phenotype upon knockdown of PP2A Cα, Aα, B56γ or SV40ST expression was determined via AI growth assay described above. To facilitate direct comparison of the MAP4K4 phosphosites across different proteomic results, all MAP4K4 phosphorylation sites were mapped and compared relative to the sites in isoform 6 of the MAP4K4 protein (O95819-6) Uniprot database (https://www.uniprot.org). Raw mass spectrometry data files are available for free download at ftp://massive.ucsd.edu/MSV000084422/.

RNAseq analysis read count was converted to Transcripts Per Million (TPM) using Kallisto quant functions (https://github.com/UCSD-CCAL/ccal) (GRCh38). Differential gene expression analysis of samples with MAP4K4 suppression vs. control was performed using mutual information. We also performed ssGSEA analysis of genesets from the literature, MsigDB (http://software.broadinstitute.org/gsea/msigdb/index.jsp), as well as IPA (https://www.qiagenbioinformatics.com/products/ingenuity-pathway-analysis) on the samples to obtain enrichment score for each genesets (Zhao et al. 2007, Barbie et al. 2009, Yu et al. 2012, Hiemer et al. 2015, Martin et al. 2018). Using the Information Coefficient (IC) (Kim et al. 2016), we estimated the degree of association of the phenotype (shMAP4K4 vs. shLuc) and their significance to the genesets.

ssGSEA and mutual information calculations: The FDRs were computed from empirical p-values using the standard Benjamini-Hochberg procedure. The empirical p-values were obtained from an empirical permutation test where the target profile is randomly permuted to generate a null distribution for the Information Coefficient (IC) values. We also generated signatures from these experiments to apply them in the CCLE RNA Seq dataset (www.broadinstitute.org/ccle) (Barretina et al. 2012) using ssGSEA. Using IC, we matched top gene dependencies associated with MAP4K4 signature score across CCLE using Gene dependency data from Project Achilles data portal using dataset version V3.12a (www.broadinstitute.org/achilles) (Aguirre et al. 2016, Meyers et al. 2017, Tsherniak et al. 2017).

Statistical analysis: All the student t-tests and p-value calculations were performed using GraphPad Prism software (https://www.graphpad.com). Unless indicated, experiments were performed in triplicates and the t-tests were performed between perturbation and relevant control conditions using triplicates values obtained from each experiment using parametric testing. Statistical significance was determined when the p-values were less than 0.05. For experiments presented in Figures 3E, 4A, 6D-E, 8, data was normalized to the mean of the controls and resulting mean values for each condition were plotted and error bars were calculated from standard deviation of the values.

## DATA AND SOFTWARE AVAILABILITY

The RNAseq data for MAP4K4 suppression experiments have been deposited in the Gene Expression Omnibus (GEO) under accession code GSE118272. Raw mass spectrometry data files are available for free download at ftp://massive.ucsd.edu/MSV000084422/.

## ACKOWLEDGEMENTS

We thank Anna Schinzel for helping with preparing lentiviral libraries. We also thank Anna Sablina and Anne-Claude Gingras for helpful discussions. LentiCRISPRv2 was a generous gift from Feng Zhang.

## Supplementary files

1. Key Resources Table

2. Normalized iTRAQ phosphoproteomic profiles of changes in phosphopetides upon suppression of PP2A Cα, Aα, B56γ or SV40ST expression.

3. Results of the SILAC experiment representing MAP4K4 interacting proteins.

4. Results of the SILAC experiment representing targeted MAP4K4 phospho-profiling.

5. Results of MudPIT experiment showing STRN4 interacting proteins.

6. RNAseq (TPM) profiles of MAP4K4 knockdown (shMAP4K4-82).

7. Genesets used in the study.

---

**Supplementary Figure 1.**
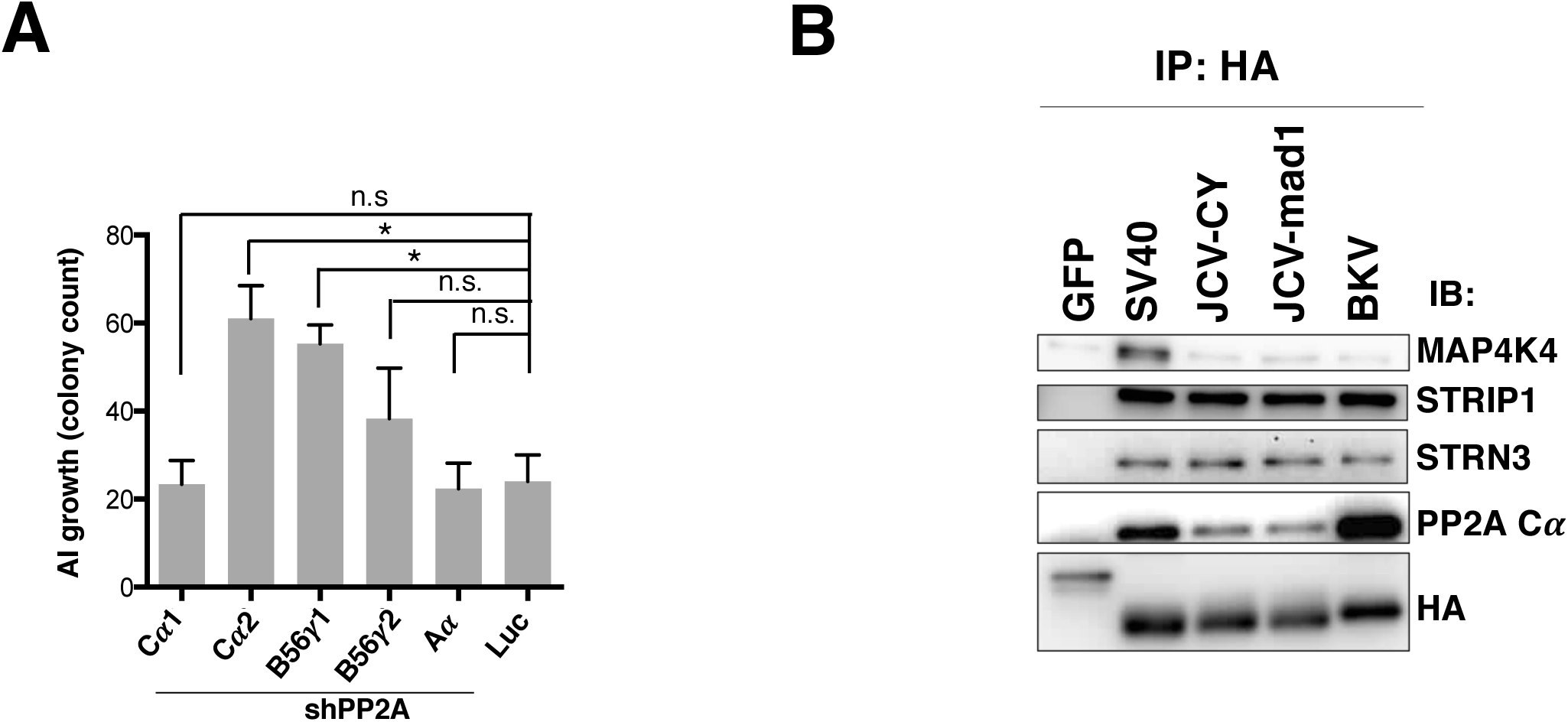
(A) AI colony count following knockdown of the indicated PP2A subunits. AI growth was assessed after PP2A subunits were suppressed using shRNAs specific for the individual subunits. (B) Interactions of polyoma virus STs with STRIPAK and MAP4K4. Co-immunoprecipitation of HA-tagged STs with components of STRIPAK and MAP4K4 confirmed results from proteomic studies (student’s t-test: n.s.= not significant, *p<0.01).

**Supplementary Figure 2.**
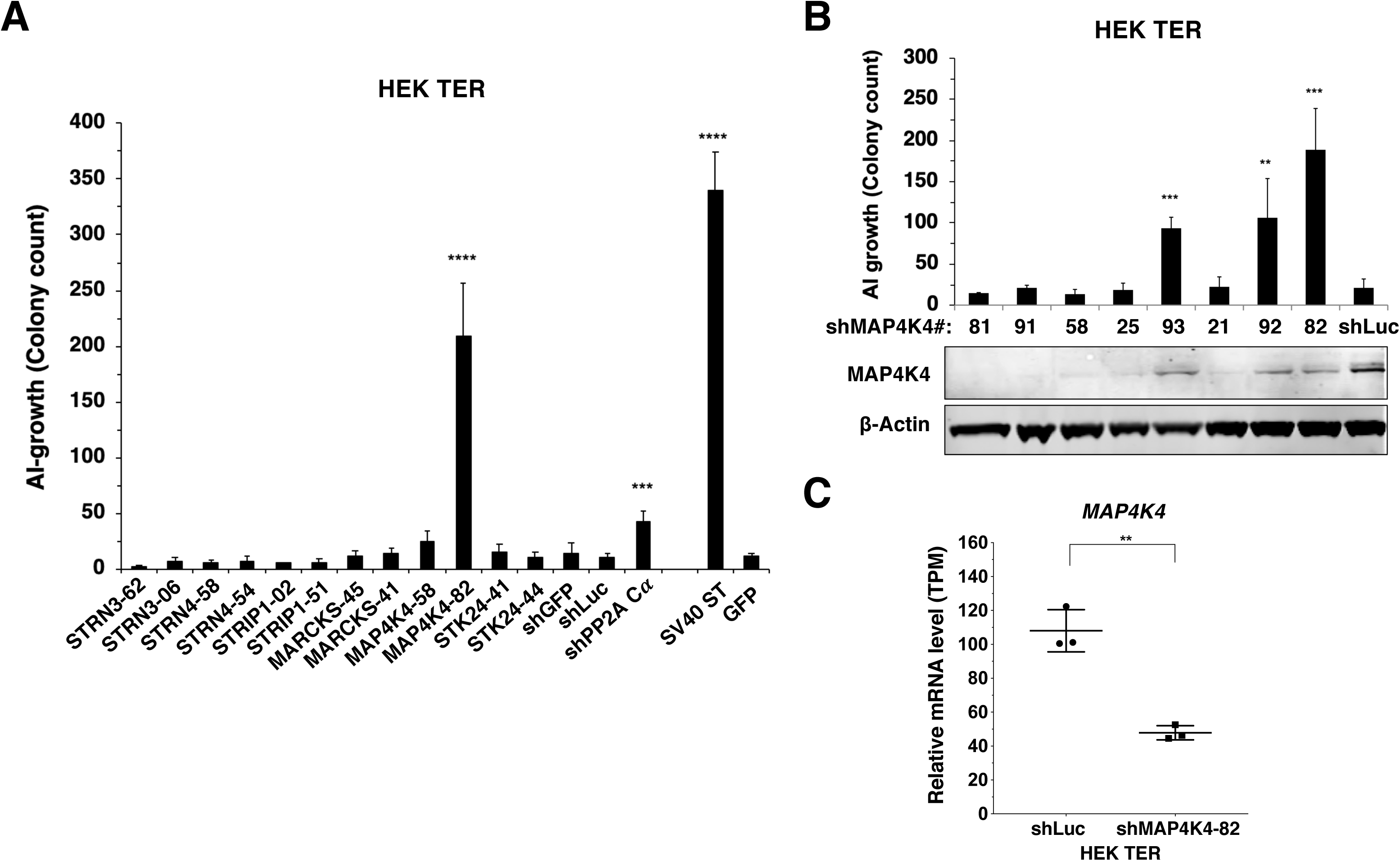
(A) AI colony count of HEK TER cells expressing the corresponding shRNAs or SV40 ST. Student’s t-test was performed for each shRNA compared to shLuc control. (B) AI colony count of HEK TER cells expressing 8 different MAP4K4 shRNAs and MAP4K4 immunoblot showing the corresponding degree of MAP4K4 knockdown. Student’s t-test was performed against shLuc control. (C) mRNA levels of *MAP4K4* after knockdown using shMAP4K4-82 as measured by RNAseq (Transcript Per Million). (student’s t-test: **p<0.001, ***p<0.0001, ****p<0.00001).

**Supplementary Figure 3.**
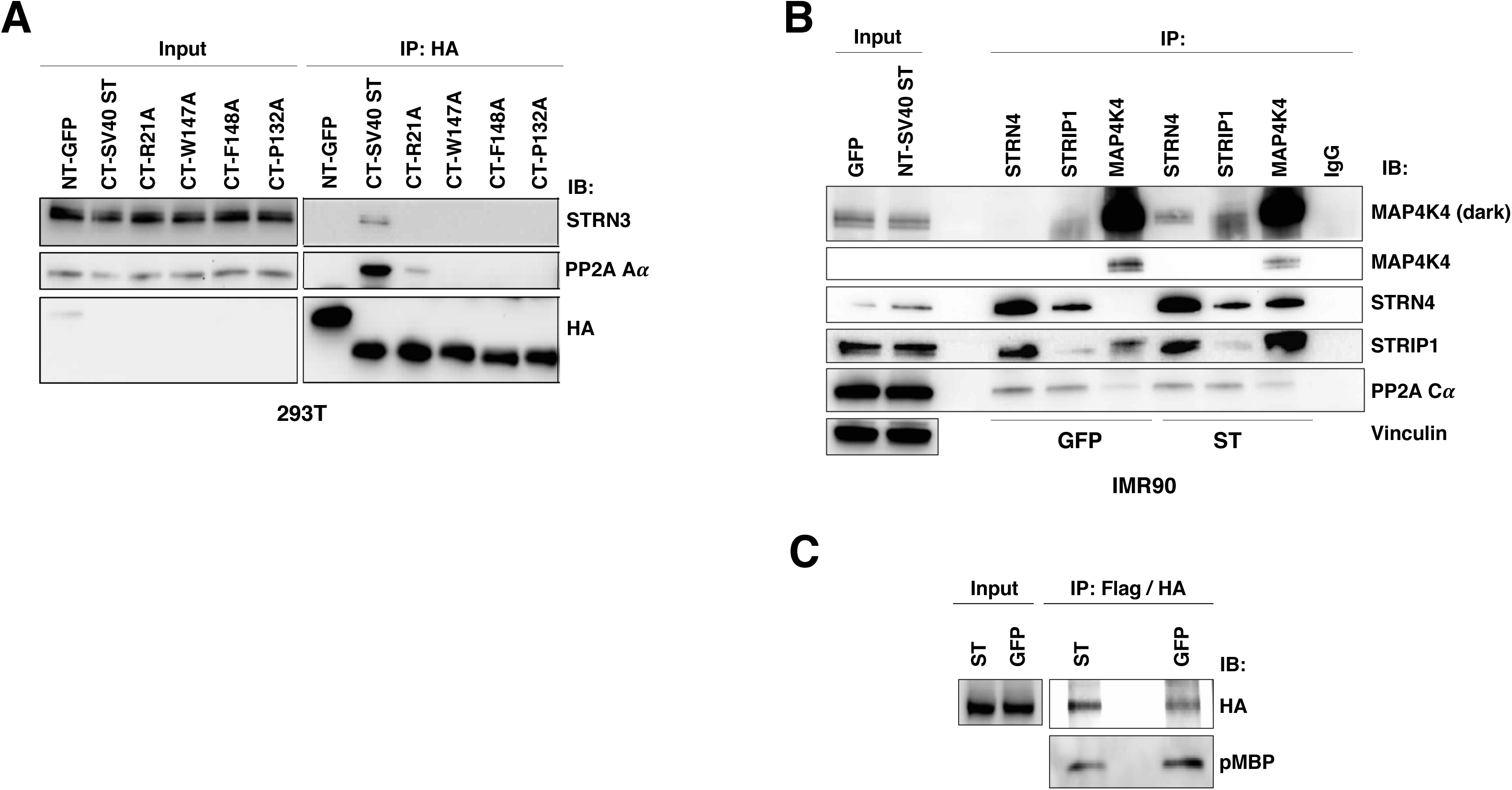
(A) Co-immunoprecipitation analysis in 293T cells of CTAP-tagged SV40 ST wild-type and PP2A binding mutants. Co-immunoprecipitation of STRN3 and A# is shown. (B) Co-immunoprecipitation analysis of STRN4, STRIP1, PP2A C# and MAP4K4 in IMR90 cells in the presence of SV40 ST or GFP. (C) *In vitro* kinase assay of tandem-affinity purified MAP4K4 (Flag/HA) from HEK TER cells expressing either ST or GFP was performed and MBP was used as a substrate. HA and phospho-MBP were detected by immunoblotting.

**Supplementary Figure 4.**
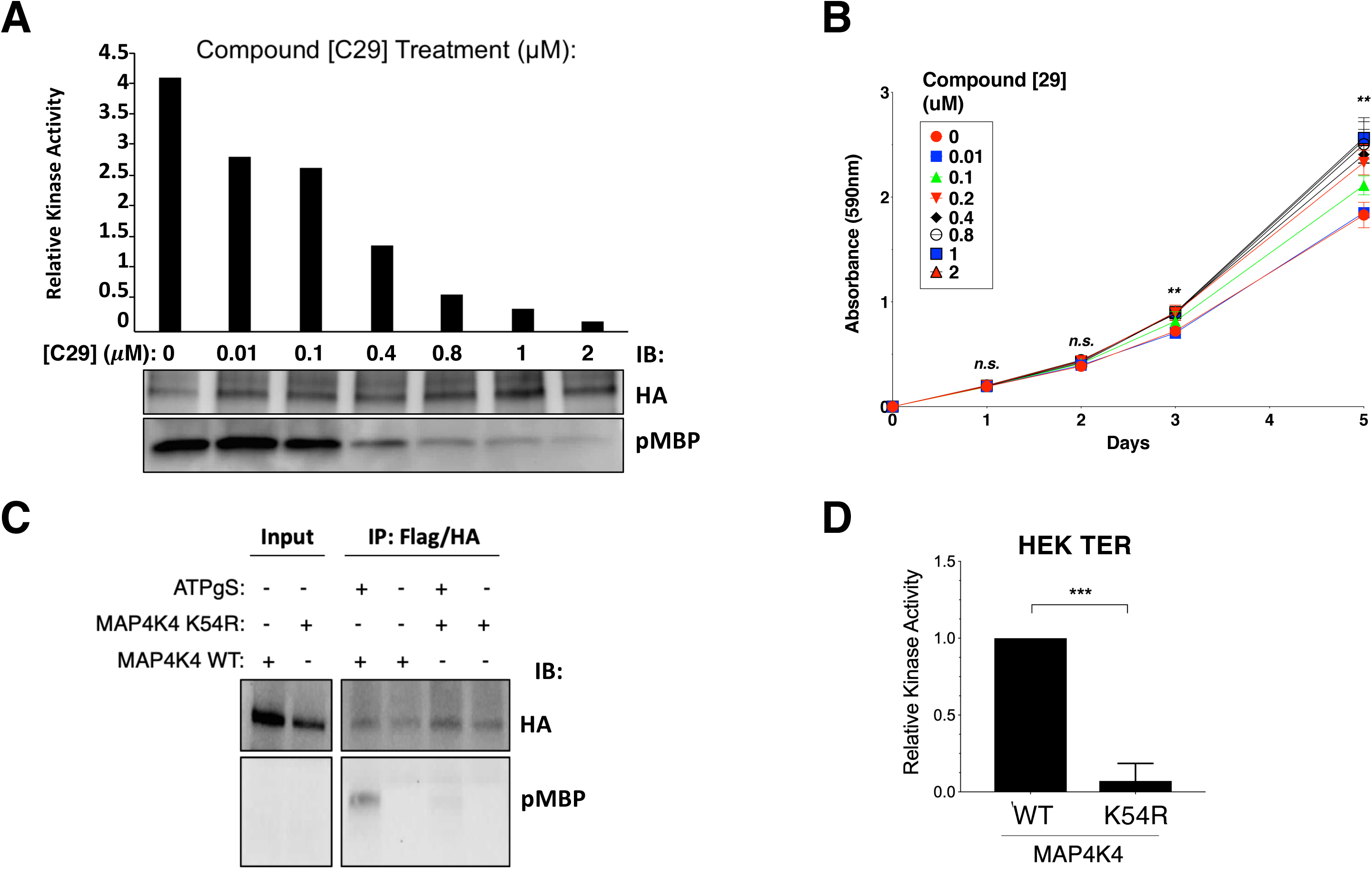
(A) *In vitro* MAP4K4 kinase assay from HEK TER cells using increasing concentrations of the MAP4K4 inhibitor (Compound 29). MBP was used as substrate and phosphorylation determined by phospho-MBP immunoblotting (bottom). Quantification of relative kinase activity (top). (B) Proliferation of HEK TER cells was measured after exposure to compound 29 at the indicated concentrations. Student’s t-test was performed based on absorbance values measured between 0 uM and 1 uM at 5 days. (C) Representative immunoblot showing MAP4K4 *in vitro* kinase assay using tandem-affinity purified wild-type (WT) or kinase-dead mutant (K54R) MAP4K4. (D) Mean MAP4K4 activity of WT relative to kinase-dead mutant (K54R) assessed using *in vitro* kinase assay (student’s t-test: n.s.*=* not significant, **p<0.001, ***p<0.0001).

**Supplementary Figure 5.**
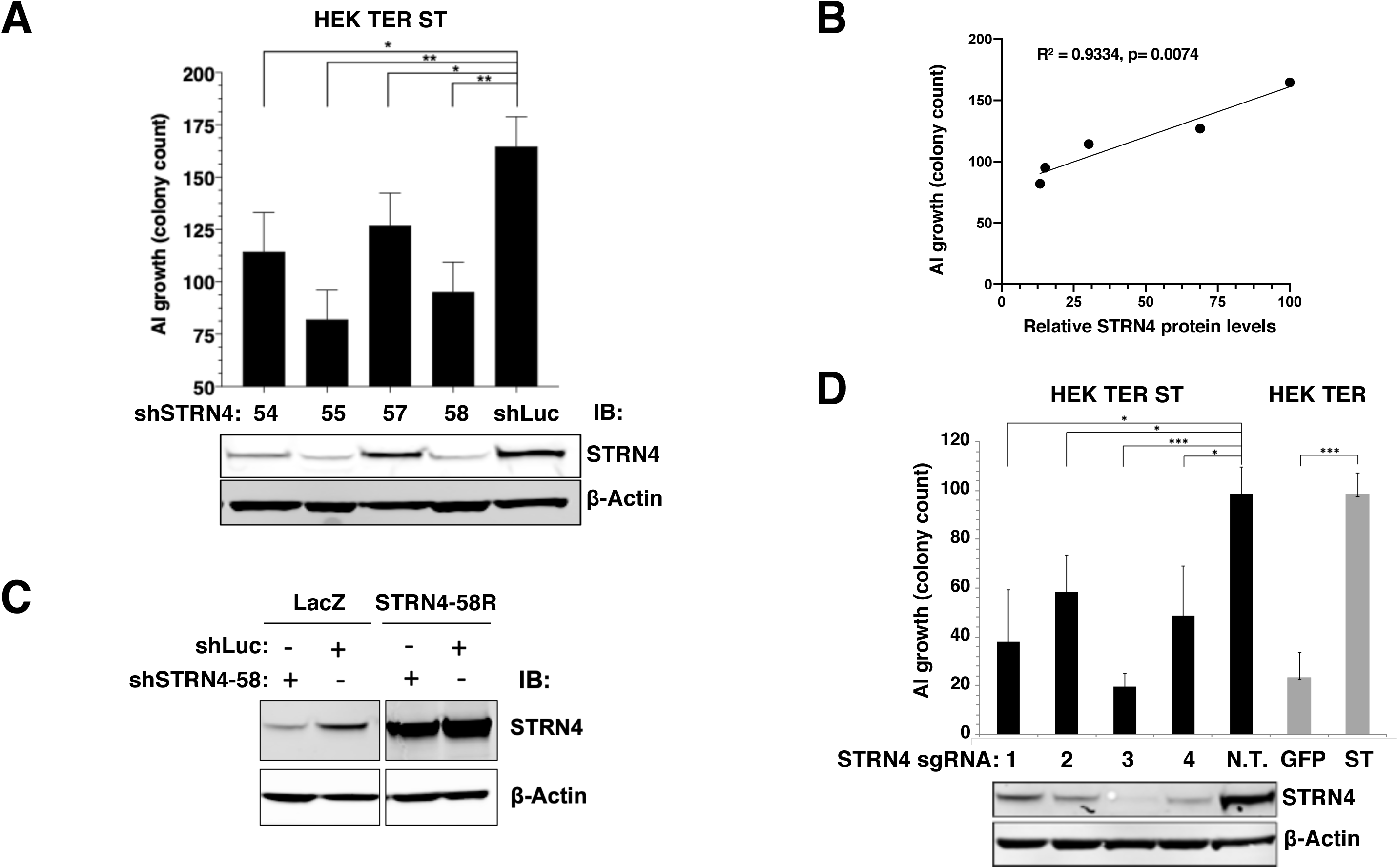
(A) AI colony count of HEK TER ST expressing 4 different STRN4 shRNAs or shLuc control. Immunoblot below depicts corresponding STRN4 expression after knockdown. (B) Correlation between the relative STRN4 protein levels (quantified and normalized to shLuc) plotted on the x-axis, and AI colony counts on the y-axis. (Pearson Correlation Coefficient and the p-values are shown). (C) Immunoblot of STRN4 in HEK TER ST cells expressing an shSTRN4-58-resistant STRN4 rescue construct (STRN4-58R) or LacZ control together with either shSTRN4-58 or shLuc control. (D) Top. AI colony count of HEK TER ST cells upon CRISPR-Cas9 mediated editing of *STRN4* using 4 different sgRNAs or a Non-Targeting (N.T.) control sgRNA. For comparison, HEK TER cells expressing ST or GFP are included. Bottom. Immunoblot depicting STRN4 protein levels after STRN4 was edited using indicated sgRNAs (student’s t-test: *p<0.01, **p<0.001, ***p<0.0001).

**Supplementary Figure 6.**
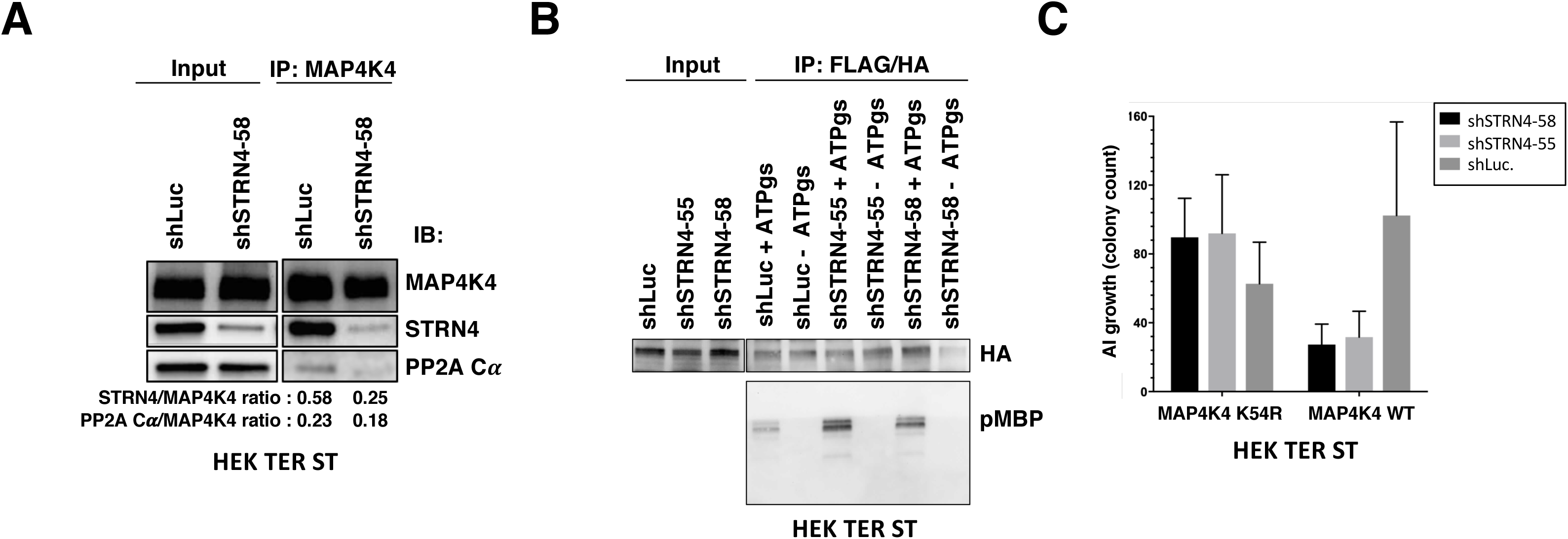
(A) Co-immunoprecipitation analysis of endogenous MAP4K4 in HEK TER ST cells with STRN4 or PP2A C” after cells were depleted for STRN4 or relative to a shLuc control. Ratio was calculated between Co-IP levels of STRN4/MAP4K4 or PP2A C”/MAP4K4 after blot quantification. (B) Representative immunoblot depicting the results of an *in vitro* kinase assay of tandem-affinity purified MAP4K4 from HEK TER ST cells expressing shSTRN4-58, shSTRN4-55 or shLuc control. (C) AI colony count of HEK TER ST cells expressing MAP4K4 kinase-dead mutant (K54R) or WT in combination with either shSTRN4-58 or shSTRN4-55.

**Supplementary Figure 7.**
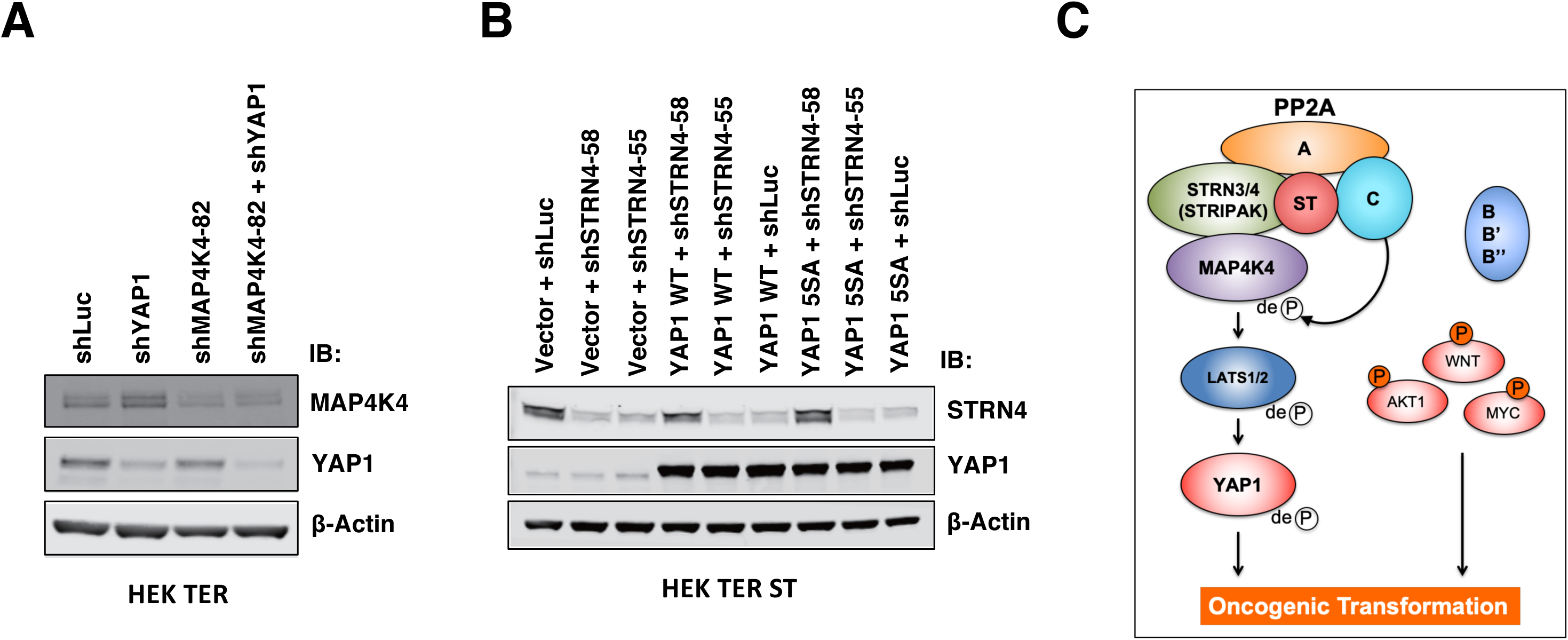
(A) Immunoblot depicting changes in MAP4K4 and YAP1 protein levels upon either depleting YAP1 alone, or in combination with MAP4K4. (B) Immunoblot depicting the overexpression of YAP1 WT or YAP1 S5A mutant, as well as suppression of STRN4. (C) Proposed model of ST-mediated transformation.

